# Mass Spectrometry Imaging of *Caenorhabditis elegans* with Fluorescence Microscopy of Tissue-Specific Reporters and Oriented Cryo-Sectioning

**DOI:** 10.1101/2025.12.01.691628

**Authors:** R. Einar Jacobson, Elizabeth Wanchisn Smith, Yasuaki Saitoh, Tian (Autumn) Qiu

## Abstract

*Caenorhabditis elegans* is a powerful model organism for studying development, neurobiology, aging, and toxicology, yet most studies rely on genetic and phenotypic measurements that do not directly capture their biochemical landscape. Mass spectrometry (MS)-based metabolomics has emerged to address this gap by enabling detection of small-molecule metabolites and lipids. Conventional chromatography-MS approaches require homogenization, eliminating spatial information during sample preparation. Mass spectrometry imaging (MSI) offers the ability to map metabolites and lipids in situ, however, its application to *C. elegans* is limited by challenges in sample preparation, particularly maintaining structural integrity during cryo-sectioning and assigning molecular features to specific tissues. In this work, we develop an integrated sample preparation workflow to overcome these constraints and enable high-resolution, spatially interpretable MSI of *C. elegans* adult hermaphrodites. We introduced a new imaging modality, fluorescence microscopy of tissue-specific, genetically encoded reporters, which expresses GFP in intestinal cells and RFP in neuronal nuclei. A mixed gelatin embedding medium preserved both structural morphology and fluorescence reporter signals during cryo-sectioning, and a simple sandwiching strategy enabled longitudinal sectioning of the worm body. Fluorescence microscopy and MSI were done on the same tissue. Using this integrated approach, we achieved matrix-assisted laser desorption/ionization (MALDI) MSI of adult *C. elegans* at 10 μm pixel resolution. By achieving structural preservation and enabling fluorescence reporters as an additional imaging modality for data interpretation, this workflow overcomes key limitations in MSI of *C. elegans* and provides a practical framework for MSI analysis of microscale biological samples.

## Introduction

*Caenorhabditis elegans*, a bacteriophagic nematode, is a widely used model animal for studying neurobiology, behavior, developmental biology, aging, and toxicology in eukaryotes.^1–4^ Its popularity arises from its ease of maintenance, tractable genetics, broad genetic homology with humans, fully documented developmental cycles for all somatic cells, and ease of observation with microscopy.^5–7^ Most *C. elegans* studies rely on measuring changes in genetics, protein expression, and phenotypic markers such as modifications to behavior, morphology, and development to study their biological processes.^8–10^ While these tools are useful for elucidating broader causal relationships between the nematode’s environment and its phenotype, they lack the ability to directly measure biological molecules with molecular specificity and interrogate biochemical mechanisms underlying phenotypic changes.

Metabolomic studies of *C. elegans* have been emerging in literature to directly measure small molecule metabolites and lipids in *C. elegans*. Particularly, mass spectrometry (MS)-based techniques have been applied to detect and annotate polar and non-polar metabolites in *C. elegans*.^11,12^ The bulk of MS-based metabolomics applied to *C. elegans* have used liquid chromatography (LC) or gas chromatography (GC)-MS to explore the *C. elegans* metabolome, where chromatography enables on-line separation of different metabolites extracted from a large population of nematodes followed by analysis by MS.^4,13–15^ While chromatography-based MS approaches provide comprehensive characterization of metabolome, a limitation of this technique is that during sample preparation worms are homogenized to extract analytes, thereby sacrificing spatial information regarding metabolite concentrations in cells and tissues.

Mass spectrometry imaging (MSI) approaches overcome the limitation by enabling acquisition of ion images regarding the *in situ* spatial distribution of metabolites and has been applied to many biological organisms. MSI workflow commonly includes cryo-sectioning of tissues, sample processing, and data analysis with mass spectrometers. Generation of biologically interpretable MSI data requires structural preservation of tissue sections and capability to localize MS features to specific tissues and cells, usually via another imaging modality such as histopathological staining. However, these two key aspects have been challenging for *C. elegans* samples due to their small size and thick cuticles. *C. elegans* is covered in a collagenous cuticle maintained by its epithelial cells which make in-situ analyte extraction difficult.^16^ Early work using freeze-and-crack helped to generate MS images but suffer from inconsistency in revealing molecular contents underneath cuticles.^17^ Thus, cryo-sectioning becomes necessary, and choices of embedding media and freezing method are important to ensure preservation of *C. elegans* structures and clean background for MSI. Previous work has demonstrated the feasibility of *C. elegans* cryo-sectioning, while structural preservation and feature spatial assignment is still challenging.^18,19^ Additionally, common imaging modalities such as histological staining that facilitate MSI interpretation are unconventional on *C. elegans* model, making it necessary to look for a different imaging modality for tissue-specific feature assignments. The challenges in structural preservation and tissue-specific assignments are intertwined, and an integrated approach is thus needed to simultaneously address these challenges.

Herein, we approached these analytical challenges by introducing fluorescence microscopy of tissue-specific fluorescence reporters as a new imaging modality and developing sample preparation strategies for cryo-sectioning strategies with controlled orientation and structural preservation. For the first time, we used tissue-specific, genetically encoded fluorescence reporters in *C. elegans* as an imaging modality on the same tissue for MSI analysis to facilitate assignment of features to tissues.^19^We utilized a dual fluorescence reporter *C. elegans* strain that expresses fluorescence proteins of different colors in intestinal cells and nucleus of neurons. We found that mixture of fish and porcine gelatin is a new embedding media that can largely maximize structural and fluorescence preservation after *C. elegans* freezing and cryo-sectioning. Using a sandwiching approach, we achieved consistent longitudinal sectioning of *C. elegans* body. With the development of sample preparation workflow, we enabled high-resolution MSI in *C. elegans* of 10-μm lateral pixel resolution using matrix-assisted laser desorption/ionization (MALDI)-MSI and revealed tissue-specific distribution of lipids and metabolites in *C. elegans* adult hermaphrodites. Our approach represents a new workflow for the MSI analysis of very small samples like *C. elegans* and adaptable for other model organisms beyond and demonstrated potential for future biological applications.

## Methods

### Materials and Reagents

Embedding media powders used were gelatin from porcine skin (PG), gelatin from cold water fish skin (FG), and sodium carboxymethyl cellulose (CMC), all purchased from MilliporeSigma. Solvents used were Optima^®^ LC-MS grade water and acetonitrile from Thermo Fisher Scientific. MALDI chemical matrices were α-cyano-4-hydroxycinnamic acid (CHCA) from MilliporeSigma and 1,5-diaminonaphthalene (DAN) from Tokyo Chemical Industries (TCI). Trifluoroacetic acid (TFA) added to CHCA solution was also purchased from MilliporeSigma. Tissue-Tek^®^ vinyl biopsy Cryomold^®^ for embedding were purchased from Sakura Finetek USA, Inc. Methanol and 2-methylbutane used for freezing were both purchased from MilliporeSigma.

### Strains and Handling

The following Caenorhabditis elegans (*C. elegans*) and bacterial strains were acquired from the Caenorhabditis Genetics Center (Minneapolis, MN, U.S.A): SJ4144 (*zcIs18 [ges-1::GFP(cyt)]*) which expresses cytosolic GFP in the intestinal cells, OH10690 (*otIs356 [rab-3p(prom1)::2xNLS::TagRFP] V*) with pan-neuronal nuclear RFP expression, and *Escherichia coli* (*E. coli*) OP50 as *C. elegans* lab food. SJ4144 and OH10690 were crossed to create the dual reporter strain (*rab-3p(prom1)::2xNLS::TagRFP, ges-1::GFP(cyt)*) used in this work. *C. elegans* were cultured on nematode growth media (NGM) plates at 20°C fed with *E. coli* OP50.

We performed bleach synchronizations on worm populations to ensure that all worms would be gravid adults during imaging. Briefly, a *C. elegans* population containing many eggs and gravid adults is rinsed off NGM culture plates using a micropipette to dispense 1 mL of M9 buffer onto a plate, then drawing the suspended worms back into the pipette. The same solution of suspended worms was used to rinse four additional plates before being deposited in a 1.7 mL centrifuge tube. Rinsing was then repeated starting from the last plate rinsed on the first pass and placed in the same centrifuge tube. Worms were centrifuged at 300 g for 1 minute before removing supernatant and replacing with 1 mL of fresh M9. This was done twice to remove bacteria, then the worms were centrifuged a final time and the supernatant removed and replaced with 700 μL of bleaching solution. Our bleaching solution was composed of 4.9 mL of ultrapure water, 0.8 mL of 6% sodium hypochlorite (Clorox) bleach, and 0.3 mL of 5 M sodium hydroxide. Worms were vortexed approximately every 30 seconds and continuously monitored under a microscope to ensure that worm bodies were almost completely dissolved. As soon as only eggs remained in solution, we quickly added 500 μL of M9 buffer to quench the bleaching before centrifuging at 1000g for 1 minute. Supernatant was replaced with 1 mL of M9 buffer before vortexing the tube and repeating the rinse two additional times to ensure that the bleach solution was fully removed. After removing supernatant for the final time, 500 μL of M9 buffer was added and the tubes placed on a rotator at 12 rpm to incubate overnight. The following day, larvae density was counted and ∼2000 L1 worms were pipetted onto 60 mm NGM plates seeded with OP50 *E. coli* as described above to grow into adults for imaging.

### *C. elegans* Embedding

Embedding media were created using powders added to water to create a gel for embedding worms. This solution was then heated in a 55 °C water bath to dissolve the embedding medium with occasional shaking or sonicating to assist in dissolution. Media were stored at 4 °C to prevent microbial growth and warmed at 30 °C until liquid before use. Four embedding media were thoroughly investigated: 20% w/v FG, 20% w/v 9:1 FG:PG, 20% w/v 19:1 FG:PG, and 5% CMC. To embed worms, we placed 350 μL of liquid embedding medium in a 10 mm x 10 mm x 5 mm cryomold. Approximately 10,000 gravid-stage *Caenorhabditis elegans* were quickly rinsed off of multiple culture plates using 150 mM ammonium formate buffer and allowed to settle to the bottom of a 1.7 mL centrifuge tube a few minutes, forming a dense worm pellet at the bottom of the tube. Worms were rinsed once to three times until no visible bacteria suspension was observed in the supernatant. Then, 50 μL of worm pellets was pipetted into the embedding medium. The worms were pipetted toward the bottom of the embedding media to prevent worm pellet from forming a liquid layer on embedding media surface. The mixture was then gently stirred with the pipette tip, which distributes worms evenly throughout the embedding media. Amongst embedding media tested, 20% FG and 5% CMC are both liquids at room temperature, while 20% 9:1 FG:PG and 20% 19:1 FG:PG are weak gels at room temperature. Therefore, 20% 9:1 and 19:1 FG:PG embedding media were heated at 30 °C into fluids and cooled at bench for several minutes before being pipetted into cryomolds for embedding.

A “sandwiching” embedding method was developed to orient the worms for cryo-sectioning at longitudinal instead of random directions. In this method, 250 μL of 20% 4:1 PG:FG stained pink with FD&C Red Dye #3 was heated to 40 °C and pipetted into a cryomold. Another 500 μL of the same embedding medium stained with FD&C Blue Dye #1 was heated and placed in a separate cryomold. After cooling until solid, the red block was cut out of the mold using a sharp craft knife and placed in a new mold such that the bottom of the block faced up to avoid placing worms on the curved meniscus surface. 20 μL of worm pellet was pipetted onto the red block and excess buffer wicked away using a clean lint-free wipe. 20 μL of embedding medium was placed on the worms before the blue block was pushed out of its mold and the bottom lowered onto the worm layer with gentle pressure applied to drive out bubbles and ensure a layer of uniform thickness. Following one of these embedding methods, molds were frozen using one of the following freezing techniques. The embedding process can be found in the “WormSandwiching movie” in the supplemental materials.

### Freezing

Three methods of freezing embedded *C. elegans* were assessed: methanol/dry ice slurry, liquid nitrogen, and isopentane, ranking from comparatively slow to fast freezing. For methanol/dry ice slurry freezing, a bath of methanol was mixed with and chilled using dry ice. An aluminum foil boat was floated on the surface of the methanol and allowed to cool and equilibrate for several minutes. Cryomolds containing embedded worms were then placed in the boat for around 6 minutes until frozen solid, at which point the entire block took on a light, opaque shade of its original color. All blocks were placed in a -80 °C freezer after freezing. For liquid nitrogen freezing, a liquid nitrogen bath was prepared with an aluminum foil boat floating on the surface in the same way as for slow methanol/dry ice freezing. After the boat equilibrated embedded worms in cryomolds were placed in the boat for around 2 minutes until visibly frozen, followed by immersion in the liquid nitrogen for a further minute to ensure full freezing. Fast freezing was accomplished by placing a beaker of isopentane in a liquid nitrogen bath. The isopentane was allowed to cool until patches of solid isopentane began to form on the bottom of the beaker, indicating that it was sufficiently cold. Cryomolds containing embedded worms were then plunged in the isopentane for 30-45 seconds before immediate removal from bath and transferring to a -80 °C freezer. These freezing procedures were performed on both unoriented and sandwiched worm sample blocks. From this point, blocks were stored at -80 °C for up to a week until cryo-sectioning and MALDI-MS imaging.

### Cryo-sectioning

A Leica CM1860 cryomicrotome (Leica Biosystems) was used to section all embedded worm sample blocks. Sample blocks were equilibrated in the cryostat for at least two hours at -20 °C, followed by cryo-sectioning at 10-μm thickness. We collected all sections by sectioning parallel to the largest face of the frozen blocks to create ∼1 cm^2^ square sections. For sandwiched blocks, the sectioning head angle was adjusted so that sections were collected parallel to the boundary between colored blocks in order to collect longitudinal sections of worms. Up to twelve sections were then placed on an indium tin oxide (ITO) coated microscope slide and thaw-mounted until the section was fully dried and there was no visible moisture on tissue to minimize delocalization and reduce re-freezing artifacts. Slides were then marked with fiducial markers for later co-registration of optical microscope images with the MALDI laser source and dried using a vacuum desiccator for at least one hour.

### Microscope Imaging

We used a Zeiss Axio Imager Z2 with Axiocam 705 camera to generate multichannel images of slides containing *C. elegans* sections. Tiling experiments were performed using Zeiss ZEN 3.6 software to collect brightfield, mCherry, and EGFP channels together and create one image of the whole slide using a 10X objective. Parameters for each channel follow. Brightfield: TL Lamp light source; 3.60 V intensity; 8 ms exposure at 70% intensity; 0.00 μm focus offset; 0 px pixel shift in x and y. mCher: Xcite Xylis lamp light source; 15% intensity; 100 ms exposure at 30% intensity; 0.00 μm focus offset; 0 px pixel shift in x and y. EGFP: identical parameters to mCher channel except with exposure time of 150 ms at 50% intensity. Zeiss fluorescence filter sets were used for fluorescence imaging, with set 38 (GFP, Ex BP 470/40; Em BP 525/50) for EGFP and Set 31 (mCherry, Ex BP 565/30, Em BP 620/60) for mCherry. Brightfield images were compressed and exported to Bruker’s FlexImaging software for co-registration with the MALDI stage, for which whole slide images were exported as .tiff files at 10% compression.

### MALDI Matrix Application

To prepare samples for MSI, we used an HTX M3+ automatic sprayer to apply MALDI chemical matrices. For positive mode MSI, we sprayed 10 mg/mL α-cyano-4-hydroxycinnamic acid (CHCA) (Sigma-Aldrich) in 70:30 ACN:H_2_O (v/v) and 0.1% trifluoroacetic acid (Thermo Scientific). Spray parameters for CHCA were 75 °C for nozzle temperature, 10 psi for nebulizer gas pressure, matrix flow rate of 120 μL/min, nozzle velocity of 1200 mm/min, track spacing of 3 mm, and 8 passes in criss-cross (CC) pattern with 0 s drying time, resulting in a matrix density of 2.67 mg/mm^2^. Sections for negative ionization mode MS imaging were sprayed with 4 passes of 10 mg/mL 1,5-diaminonaphthalene (DAN) (Tokyo Chemical Industries) in 70% acetonitrile. Spray parameters for DAN were identical to those of CHCA but with a track spacing of 2 mm rather than 3 mm, resulting in a matrix density of 2 mg/mm^2^. MSI was performed on slides immediately after matrix application.

### MALDI Mass Spectrometry Imaging

MS images were obtained using a Bruker timsTOF fleX mass spectrometer. Calibration was performed in ESI mode using a 10 mM sodium formate (12.5 mL water, 12.5 mL isopropanol, 50 μL concentrated formic acid, and 250 μL 1 M NaOH) to add calibrant peaks in the low mass range. Enhanced quadratic calibration was used and accepted when calibration score was reported as greater than 99% by timsControl software. Online calibration was performed using MALDI matrix ion peaks as lock masses. The *m/z* used were 212.032 for CHCA matrix ions and 155.062, 156.069, 157.077, 313.046, and 471.230 for DAN matrix ions.^20^ After data acquisition, matrix ions were re-examined to ensure they are within 10 ppm of mass error compared to their exact masses. For datasets that experience systematic mass shift over 10 ppm, a recalibration process was performed using Bruker’s DataAnalysis software based on matrix ions.

For data acquisition, a Bruker MTP Slide-Adapter II was used to position two slides in the MALDI source. For positive mode with CHCA, masses from 20-650 *m/z* were analyzed with 10 μm single beam-scan square laser pattern with laser intensity of 35%, frequency of 10 kHz, and 200 shots per pixel. Ion optics parameters were as follows: MALDI plate offset of 50 V; deflection 1 delta of 60 V; funnel 1 RF of 300 Vpp; isCID energy 0 eV; funnel 2 RF 200 Vpp; multipole RF 200 Vpp; quadrupole ion energy 10 eV; low mass cutoff 20 *m/z*; pre-TOF transfer time 55 μs and pre pulse storage 3 μs. This method provides an average mass resolution of 36663 at 471.146 *m/z* and mass accuracy of 1.4 ppm for positive mode with CHCA. Average mass accuracy was calculated based on known matrix ion cluster masses at 190.050, 212.032, and 221.972 for CHCA.

For negative mode with DAN, mass range was extended to 100-1050 *m/z* to enable lipid analysis. Laser intensity was maintained at 35% with 10 kHz repetition rate and 150 shots per 10-μm laser beam scan. Ion optics parameters were as follows: MALDI plate offset of 50 V; deflection 1 delta of -70 V; funnel 1 RF of 350 Vpp; isCID energy 0 eV; funnel 2 RF 200 Vpp; multipole RF 200 Vpp; quadrupole ion energy 5 eV; low mass cutoff 50 *m/z*; pre-TOF transfer time 50 μs and pre pulse storage 5 μs. This method provides an average mass resolution of 25405 at 221.972 *m/z* and mass accuracy of 1.2 ppm for negative mode with DAN. Average mass accuracy was calculated based on known matrix ion cluster masses at 156.069, 157.077, 313.046, and 471.230 for DAN. Mass resolution data for all matrix ion clusters is available in **Table S1**.

### Data Analysis

MS imaging data analysis was done using Bruker’s SCiLS Pro Software. Datasets were total ion current (TIC) normalized before identifying features associated with *C. elegans*. Worm feature lists were created in Bruker’s SCiLS Lab software by importing all features with a 0.3% relative intensity threshold. These features were then manually searched and only signals localized to *C. elegans* bodies were kept. Co-localization of features between the worm region and a larger region that contains worm and adjacent areas was examined using the “Find values co-localized to region” function in SCiLS, which returned correlation scores for each feature with higher scores indicating greater co-localization of signals to worm bodies.^21,22^ MS1 features were used to manually search for matches in LIPID MAPS, Human Metabolome Database (HMDB), and Metabolomics Workbench databases for putative annotation. Mass lists were uploaded to each database either by copying lists or uploading a text file. All searches were conducted with a 0.005 *m/z* tolerance. Data were also exported from SCiLS as .mzML and .ibd file types for upload to METASPACE and searched at 10 ppm mass tolerance using the METASPACE-ML analysis version to search the CoreMetabolome – v3, ChEBI – 2018-01, HMDB – v4, and LipidMaps – 2017-12-12 databases with 5% and 10% false discovery rate (FDR) levels. A manually curated, *C. elegans*-specific metabolome database was also examined for possible matches.^23^

## Results

### Testing various embedding media and freezing methods for *C. elegans* cryo-sectioning

To improve the preservation of *C. elegans* anatomy, we first tested various embedding media as well as three different freezing methods by mixing worm population with embedding media followed by freezing (**Figure 1**). Initial sectioning trials by using a solution of 20% w/v gelatin from fish skin (fish gelatin) enabled the identification of some structures but exhibited damage to worm bodies. We hypothesized that this damage was incurred during sectioning, indicating the need to use an embedding medium with higher gel strength to support the worm bodies and prevent their deformation. Fish gelatin at 20% is fluid at room temperature. Gelatin derived from mammalian tissue has been reported as a useful embedding medium with higher gel strength than fish gelatin, but trial solutions of 5-20% w/v porcine gelatin demonstrated a gel point too high to avoid heat shocking or outright killing embedded worms.^24–26^ To achieve a balance between the need for sufficient gel strength and low gelling point, we created mixtures of fish gelatin and porcine gelatin and decided a 20% w/v 9:1 and 19:1 fish:porcine gelatin mixture should be further tested for embedding due as these compositions provide a higher gel strength than 20% fish gelatin, but at a gel point of ∼27 °C. To minimize heat shock during embedding, gel was heated to fluid in 30°C bath and briefly cooled on bench before rapid mixing with worm populations and subsequent freezing. After initial optimization, four embedding media were chosen for further investigation, including 20% fish gelatin, 20% w/v 9:1 and 19:1 fish:porcine gelatin mixture, as well as 5% carboxymethylcellulose (CMC), an embedding medium which has been reported effective for other small animals such as *Danio rerio* embryos and *Daphnia magna*.^27–29^ Furthermore, four freezing methods were investigated, including a dry ice/methanol slurry, vapor freezing over liquid nitrogen before immersion, and direct immersion in isopentane cooled by liquid nitrogen, as described in Method section. These methods freeze embedding blocks at different rates, with dry ice/methanol being the slowest and isopentane freezing being the fastest.

**Figure 1.**
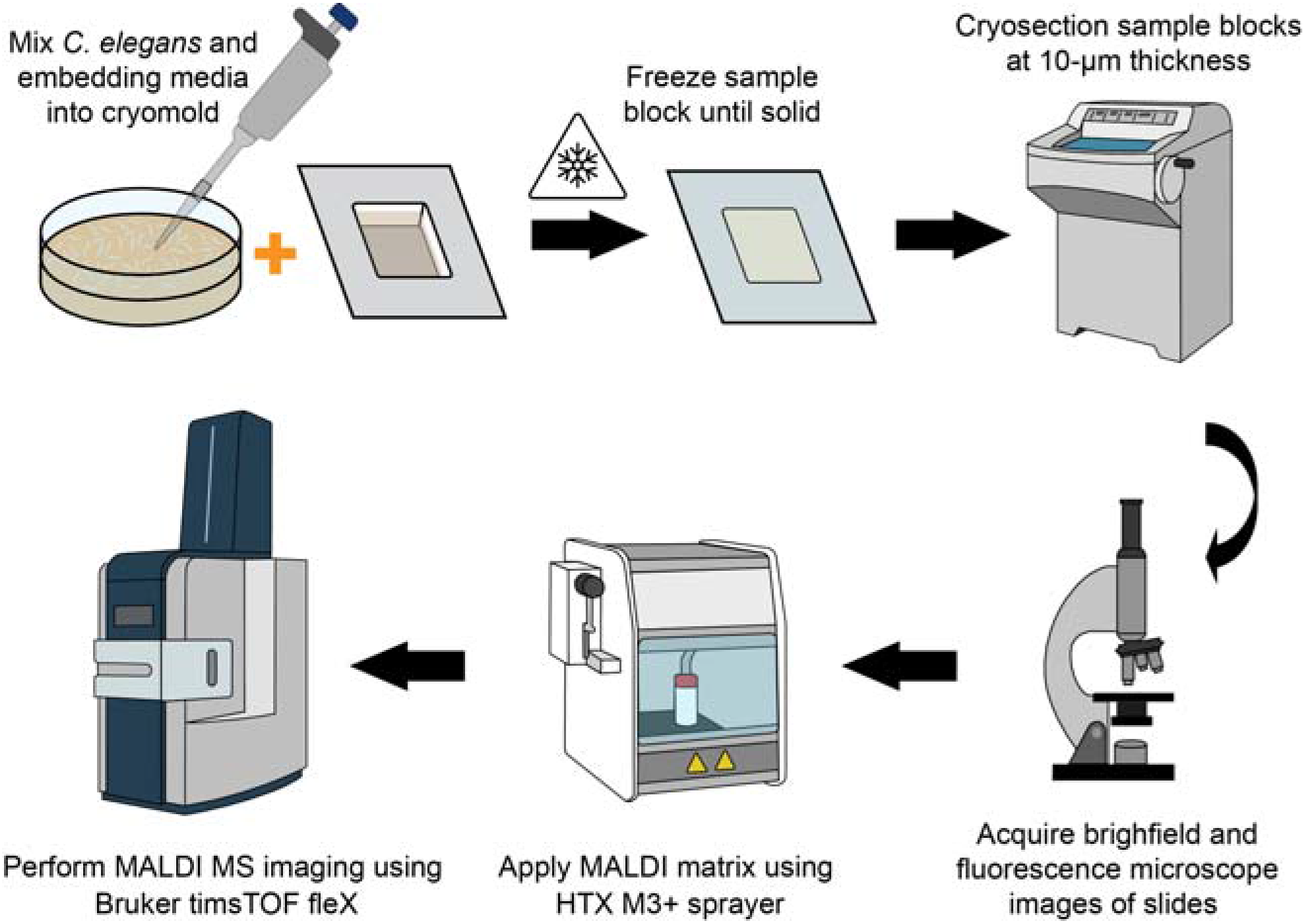
A flowchart of the steps used in preparing *C. elegans* for MALDI-MS imaging with unoriented embedding. Adult worms were rinsed off of culture plates using ammonium formate solution. Worms were pipetted into a centrifuge tube and allowed to settle before pipetting and gently mixing them into embedding medium which had been placed in a cryomold. Molds were then frozen and stored at -80 °C until sectioning. Sections were collected at -20 °C and 10-μm thickness after equilibrating blocks for 2 hrs. Sections were thaw mounted on ITO slides and vacuum desiccated before brightfield and fluorescence images collected using a Zeiss Axio Imager Z2. MALDI chemical matrices were then applied to slides using an HTX M3+ pneumatic sprayer before performing MALDI-MS imaging with a Bruker timsTOF fleX.

The results of these trials are shown in **Figure 2**. We categorized worm sections into two orientations for assessment. Longitudinal sections are those in which the worm was cut along a significant part of its length resulting in an oblong section, while cross sections cut across the worm’s width for roughly circular sections. Worm sections were mostly cross sections with a small portion of longitudinal sections, due to the shape of worm from unoriented mixing and embedding. We observed that 5% CMC was an inadequate embedding medium as both longitudinal and cross sections were severely damaged with an “exploded” look, in which no anatomical structures can be identified (**Figure 2A-A’’**). Unsurprisingly, MS images also showed severe delocalization for CMC. The gelatin-based embedding media performed better in worm structure preservation. For 20% fish gelatin, cross sections were kept mostly intact, while longitudinal sections were often severely damaged, leaving “gaps” between cuticle and tissues (**Figure 2B-B’’**). Both 20% 9:1 and 19:1 fish:porcine gelatin performed better on longitudinal sections while preserving cross sections, but the higher gel strength of 9:1 F:P gelatin allowed more reliable observation of intact longitudinal sections across different freezing methods (**Figure 2C-C’’, D-D’’**).

**Figure 2.**
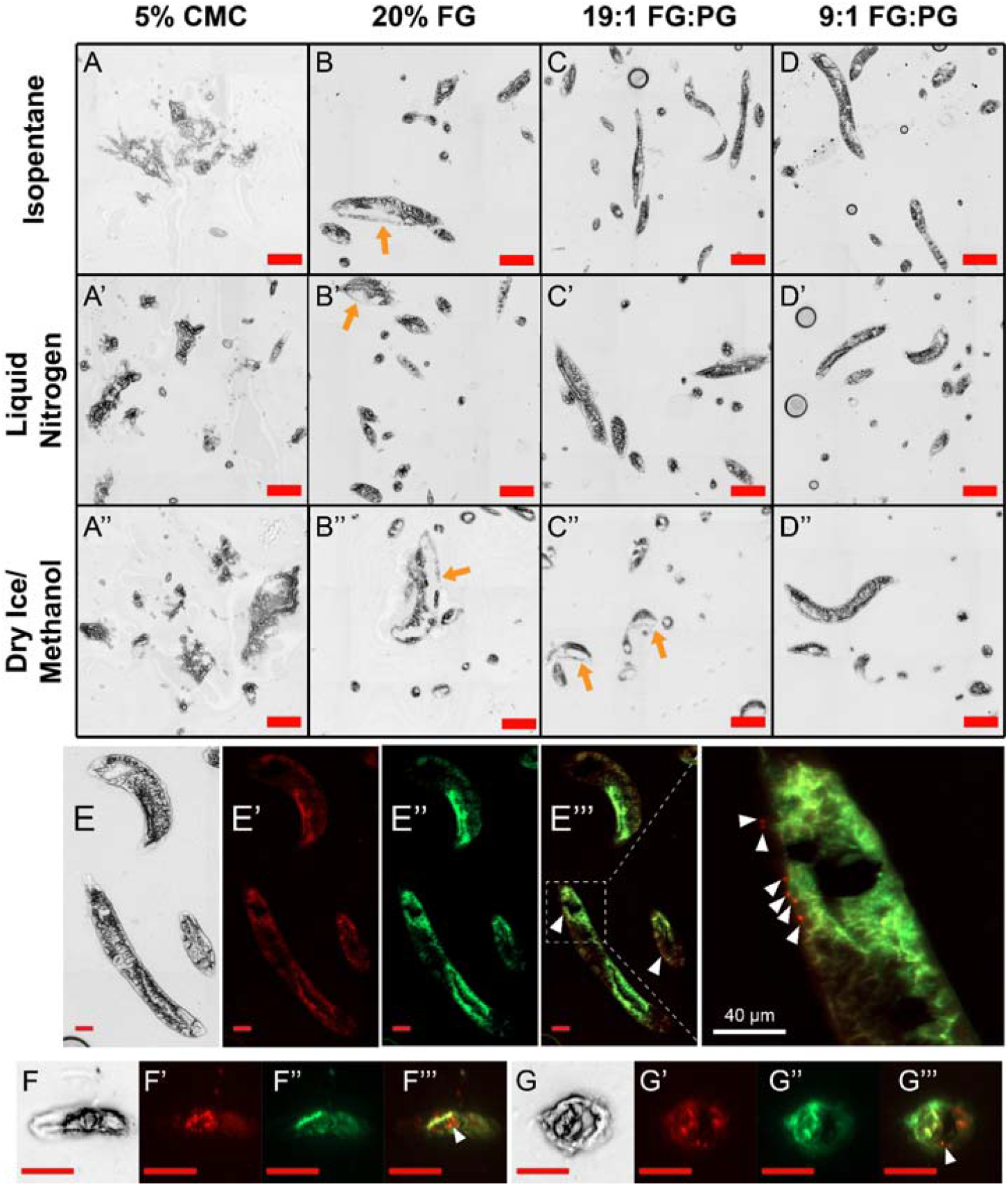
Panel A-D’’ depicts regions of *C. elegans* with different freezing methods and embedding media. From left to right the media are 5% CMC, 20% FG, 19:1 FG:PG, and 9:1 FG:PG. From top to bottom the freezing methods are liquid nitrogen cooled isopentane immersion, liquid nitrogen freezing, and freezing over dry ice/methanol bath. Orange arrows point to the “gaps” between cuticle and tissue as a manifestation of tissue damage. Panel E-G’’’ shows brightfield, RFP fluorescence, GFP fluorescence, and merged fluorescence images of representative *C. elegans* sections embedded in 9:1 FG:PG frozen using liquid nitrogen. Panel E-E’’’ display longitudinal partial whole-body sections. Panel F-F’’’ display a worm head. Panel G-G’’’ display a cross section through a worm midbody. White triangle indicates nuclei of neurons with RFP. Scale bars represent 200 μm for A-D’’ and 50 μm for E-G’’’ unless otherwise marked.

The role of freezing method on worm structure preservation appears to be less important than embedding media. For 20% fish gelatin only and 9:1 FG:PG, freezing methods did not generate an observable effect in worm structure preservation (**Figure 2B-B’’, 2D-D’’**). However, for 19:1 FG:PG, it seemed that dry ice/methanol performed less optimal than liquid nitrogen and isopentane by assessing the damage in longitudinal sections (**Figure 2C-C’’**). For 5% CMC, altering freezing method did not rescue the structural damage (**Figure 2A-A’’**), indicating that the embedding media cannot hold the worm bodies intact during cryo-sectioning. To explore further enhanced protection of worm bodies from freezing and cryo-sectioning, we tested cryo-protectant solutions such as sucrose and glycerol by doping them into the viscous embedding media, as the use of cryo-protectant solutions is a common practice in cell viability preservation during cryo-storage.^30^ We tested doping 2 M sucrose as well as 1.5 M glycerol into each.^30–32^ We found that the addition of cryoprotectant depresses the melting point of the embedding medium to the extent that cryo-sectioning became not viable, leading us to abandon this approach in favor of a simpler optimized embedding and freezing protocol.

### Examining preservation of cell-specific fluorescence markers after cryo-sectioning

Tagging tissue-specific marker genes with fluorescence protein reporters are widely used in *C. elegans* research to visualize certain tissues and subcellular compartments through the transparent body of worms.^33^ The genetic encoding is tractable and can be multiplexed. To explore the possibility of using genetically encoded fluorescence reporter as a new imaging modality to align *C. elegans* MSI data, we crossed two reporter strains available from CGC and created a dual reporter strain with intestine cytosolic GFP and pan-neuronal nuclear RFP (*rab-3p(prom1)::2xNLS::TagRFP, ges-1::GFP(cyt)*) in this work. **Figure S3** showed whole live worm images, which showed clear GFP in intestinal cells and RFP signals as reporters in neuronal nuclei and as autofluorescence in intestinal cells originated from gut granules, consistent with the known intestinal autofluorescence in *C. elegans*.

We evaluated the preservation of fluorescence reporter signals after cryo-sectioning via microscopy. *C. elegans* were embedded in three gelatin-based embedding media; the 5% CMC media was dropped because of the severe structural damage observed from brightfield images. Both cross sections of worm head and mid-body (**Figure 2F-G’’’, Figure S1**) and longitudinal sections (**Figure 2E-E’’’, Figure S2**) were examined. In the fluorescence images of sectioned *C. elegans*, we observed strong fluorescence signals from the intestinal cytosol GFP that co-localize with the autofluorescence in RFP channels. The RFP channels showed generally high autofluorescence, making it challenging to differentiate neurons solely from red channels (**Figure 2E’, F’, G’**). While locating neurons can be challenging in RFP only images, merging the RFP channel with the GFP channel quickly reveals neurons as clear red dots, as the red autofluorescence overlap with cytosol GFP signals to take on a yellowish hue, making the highly localized nuclei-specific RFP signals stand out (**Figure 2E’’’, F’’’, G’’’**). We also noticed that there was non-specific fluorescence that did not seem to localize to intestine but co-localize to high-contrast areas with brightfield images (e.g., **Figure 2F’’ vs F**). We suspect such non-specific fluorescence signals were from diffusion or scattering of uneven surface after sectioning. In general, autofluorescence and non-specific fluorescence across the three embedding media and freezing methods were generally comparable (**Figure S1, S2**), with some variations from trial to trial but not consistently higher/lower from any of the embedding media or freezing methods. Therefore, liquid nitrogen freezing with 9:1 F:P gelatin embedding was chosen for use in subsequent MALDI-MS imaging due to its success in preserving both worm structure and fluorescence reporter signals, as well as its ease in implementation and lower fire hazard.

### Mapping metabolites in *C. elegans* cross sections to fluorescence reporter-expressing tissues

Using worms embedded in 20% 9:1 FG:PG frozen by liquid nitrogen, we performed MALDI-MSI on worm sections from unoriented embedding with a focus on cross sections. Most worm sections embedded in this way are cross sections of mid body, with a few cross sections of worm heads and partial longitudinal sections also observed. Fluorescence and selected MS images of a few representative worm sections are shown below in **Figure 3**. MS images collected using DAN are shown in panels A and B, with those collected using CHCA shown in panel C. Representative average spectra of unoriented sections are shown in **Figure S4 and S5** (top panels). Putative annotations based on *m/z* of shown ion images are listed in **Table S2**, while it should be noted that these annotations are only based on MS1 with limited confidence of identification. DAN was selected for negative mode analysis due to its basicity and excellent performance in negative mode, and CHCA was used as a classical MALDI matrix for positive mode analysis.^34^ Both DAN and CHCA yield a uniform chemical matrix layer comprised of small crystals when applied using the aforementioned sprayer settings. Ideally, a sublimation apparatus provides the smallest crystal size, however, the unavailability for sublimation apparatus in our laboratory limits the testing of sublimation preparation, which is a future research direction. Lateral (pixel) resolution and laser spot size of 10 μm was chosen to balance the needs between detection sensitivity and high-spatial resolution due to the small size of *C. elegans*.

**Figure 3:**
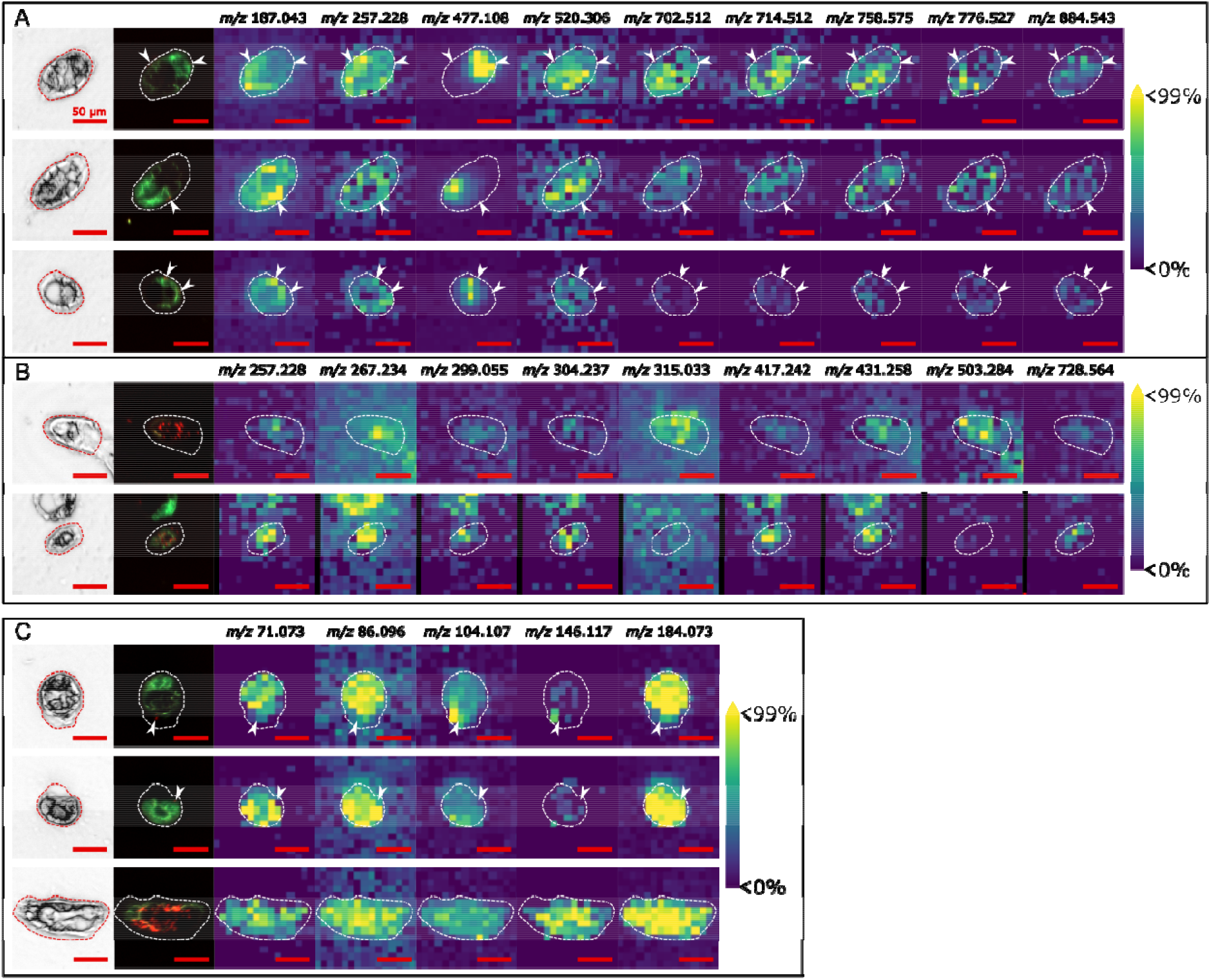
Cross-sections of *C. elegans* imaged using DAN and CHCA MALDI chemical matrices. Panel Set A and B are a collection of cross-sectional images using DAN as a chemical matrix, with midbody sections in Panel Set A and sections containing neural ganglia in Panel Set B. Panel Set C contains images collected using CHCA as MALDI matrix, with the first two rows showing midbody sections and the bottom row showing a section through the worm’s head (note that this head section appears to have been sectioned with an angle, showing an oblong shape. Dashed outlines indicate the shape of worm sections based on optical images. Scale bars are 50 μm.

In general, we observed that ions are largely localized within the tissue edges, indicating that the selected embedding media and freezing method successfully preserved the localization of metabolites during the sample preparation. **Figure 3** demonstrated spatial distribution patterns of some representative ion signals from negative mode MSI. We quantified the extent of ion signal delocalization by calculating the co-localization correlation values between a region of interest (ROI) outlining worm tissue edge and a ROI containing the worm section and nearby areas (**Figure S6**).^21,22^ Results showed that most of the ions displayed high co-localization, indicating that the ions are generally localized within worm tissues (**Table S3**). By comparing ion images to fluorescence markers with an understanding of *C. elegans* anatomy, we were able to infer the location of specific ions. **Figure 3A** shows three mid-body cross sections, determined by the intestinal GFP on one side of the section and the presence of a single neuron, presumably from the ventral nerve cord (VNC). Ion images showed that ion signals are localized differently in the same section. For example, comparing ion images to fluorescence images in three different sections, *m/z* 187.043 seemingly localized with regions of near intestinal cells, potentially gonad and peripheral area, while ion of *m/z* 477.108 strongly co-localized with fluorescence signals from intestinal cells. **Figure 3B** shows cross sections with cluster of neuronal RFP signals, indicating that these cross sections were from head or tail ganglia. Interestingly, we noticed that ion of *m/z* 257.228 was localized to at least part of the ganglia in head/tail sections, but relatively more uniformly distributed across the mid-body sections. Other ganglia-localized signals were identified, including *m/z* 267.234, 299.055, 304.237, 315.026, 417.242 and more, while their intensities vary between two different sections. Considering the observation of GFP signal in the section in the bottom row, it is possible that the top section in **Figure 3B** was from ganglia and the bottom section was from tail, which cannot be differentiated without further evidence.

**Figure 3C** presents ion images from positive mode MSI for two midbody sections (top two row) and a head section (bottom row), determined by the fluorescence reporters. Ion signals of *m/z* 71.073, 86.096, and 184.073 roughly co-localize across all three sections. They are likely the precursor and fragment ions of phosphocholine (PC) headgroup because of their mass match and co-localization. Fragmentation of PC headgroup to the same *m/z* from collision induced dissociation were reported before, whereas the fragmentation from MALDI-MSI was likely due to MALDI-induced in-source fragmentation.^35^ Ion of *m/z*

146.117 is highly localized in several pixels in the mid-body sections, co-localizing to the position of VNC neuron, and abundant in the head ganglia section, and its mass matches the monoisotopic mass of acetylcholine [M]^+^ ions. *C. elegans* possess cholinergic neurons in both VNC and head/tail ganglia, further increasing confidence in the assignment of *m/z* 146.117 to acetylcholine.^36^ Ion of *m/z* 104.107 matches the mass of choline [M]^+^ ion. Choline is a biosynthetic precursor for acetylcholine, while it can also be a fragment from PC headgroup. Therefore, it was not surprising to observe that the spatial distribution of *m/z* 104.107 was seemingly a combination of the patterns from both *m/z* 146.117 and 184.073.

As this MS imaging method had demonstrated its ability to reveal the spatial localization of ion signals, the assignment of ion signals to worm tissue was still limited due to the use of random cross sections, even with the assistance of fluorescence reporters. We thus decided to further develop an embedding method to enable controlling of worm orientation for longitudinal sections from head to tail, aiming to better analyze the tissue-specific distribution of metabolites mapped to worm anatomy.

### Achieving longitudinal sectioning and MALDI-MSI with “sandwiched” embedding

We controlled the orientation of *C. elegans* using a sandwiching method as described in Method section and pictured in **Figure 4**. Pressing worms and embedding medium between two solid blocks of gelatin (20% 4:1 porcine:fish gelatin) at room temperature ensured that they were alive until freezing while also restricting their orientation in two dimensions. During cryo-sectioning, the cutting direction was guided via the boundary between two different colors of “sandwiching” blocks to enable longitudinal sectioning of worms embedded between the blocks. Initial testing between 20% 9:1 and 19:1 FG:PG as embedding medium between the two blocks showed that 9:1 FG:PG generated more consistent results, consistent with results from unoriented embedding and sectioning. Compared to the sections from unoriented embedding that generated mostly cross sections, the sandwiched embedding and guided sectioning resulted in mostly longitudinal sections (**Figure S7**). Worm intestine is clearly visible in either complete or partial longitudinal sections, allowing us to assign ion localization to worm structures. In some sections, multiple neurons in head ganglia (**Figure 5, Section 1; Figure S8**) or VNC along one side of the worm (**Figure 5, Section 2; Figure S8**) can be observed. We did notice that perfect head-to-tail longitudinal sections were still rare, potentially due to the cylindrical but tapered worm body shape and the imperfection of alignment of worms between the two blocks. By examining serial sections of sandwiched worms, the different section planes of the same worms were clearly visible (**Figure S9**). Similar observations were observed in a recent report that used a microfluidic device to trap and align worms.^19^ Despite that, we were still able to observe good preservation of worm structure and fluorescence reporters after sandwiched embedding and oriented sectioning.

**Figure 4.**
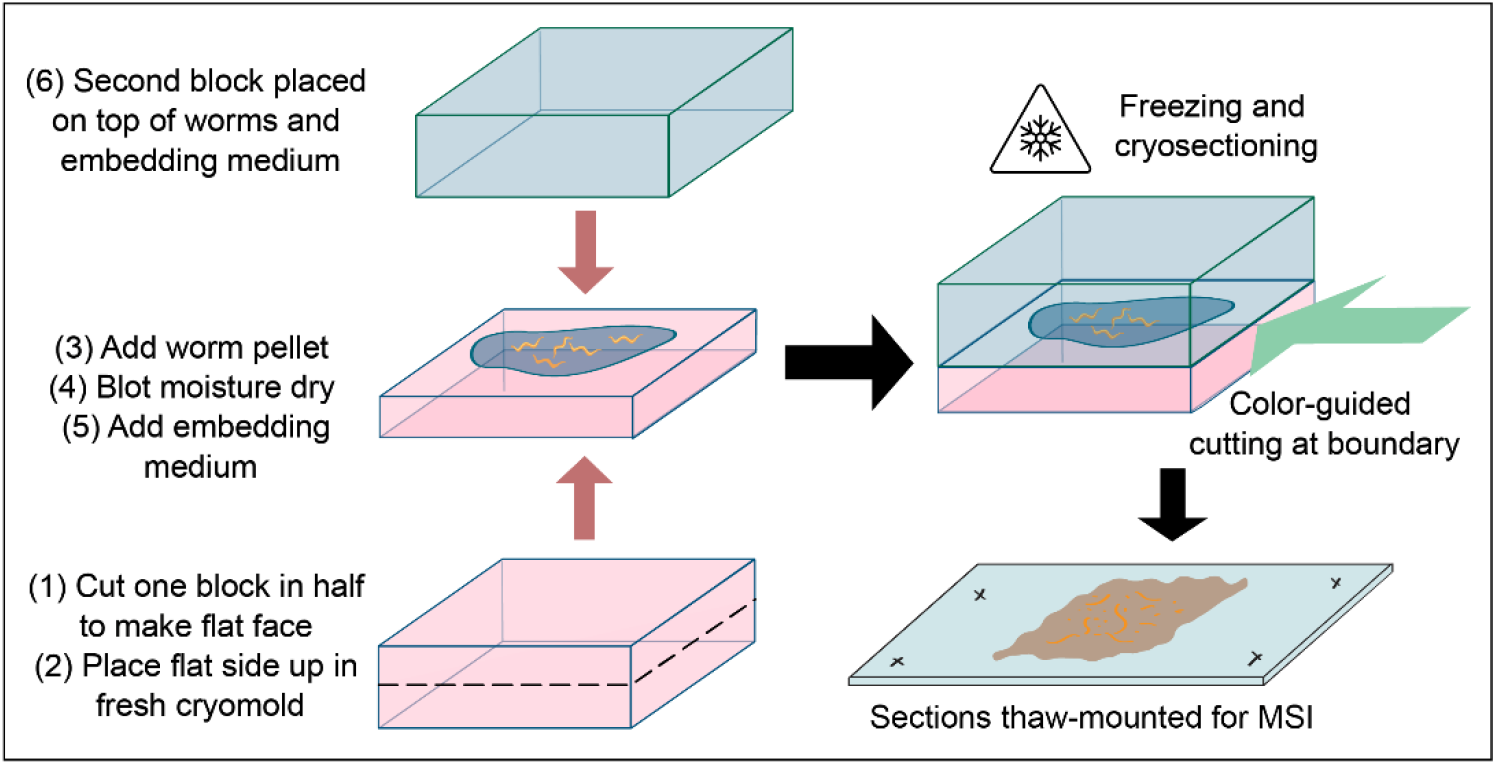
A figure outlining the “sandwiching” approach for achieving longitudinal *C. elegans* sections. Two blocks of 4:1 porcine:fish gelatin lightly colored with FD&C Red #3 and Blue #1 were formed using the same cryomolds from unoriented sectioning. One of these was removed from the cryomold and cut in half parallel to its square face. The half with the smoother side was then placed smooth side up in the bottom of a fresh cryomold to create a flat surface on which to place worms for embedding. 20 μL each of gravid worm pellet and embedding medium were pipetted onto this surface. The second block was then placed on top of the worms and embedding medium, using a pipette tip to press it down such that any trapped air could escape. This assembly was then frozen using liquid nitrogen vapor as described previously.

**Figure 5.**
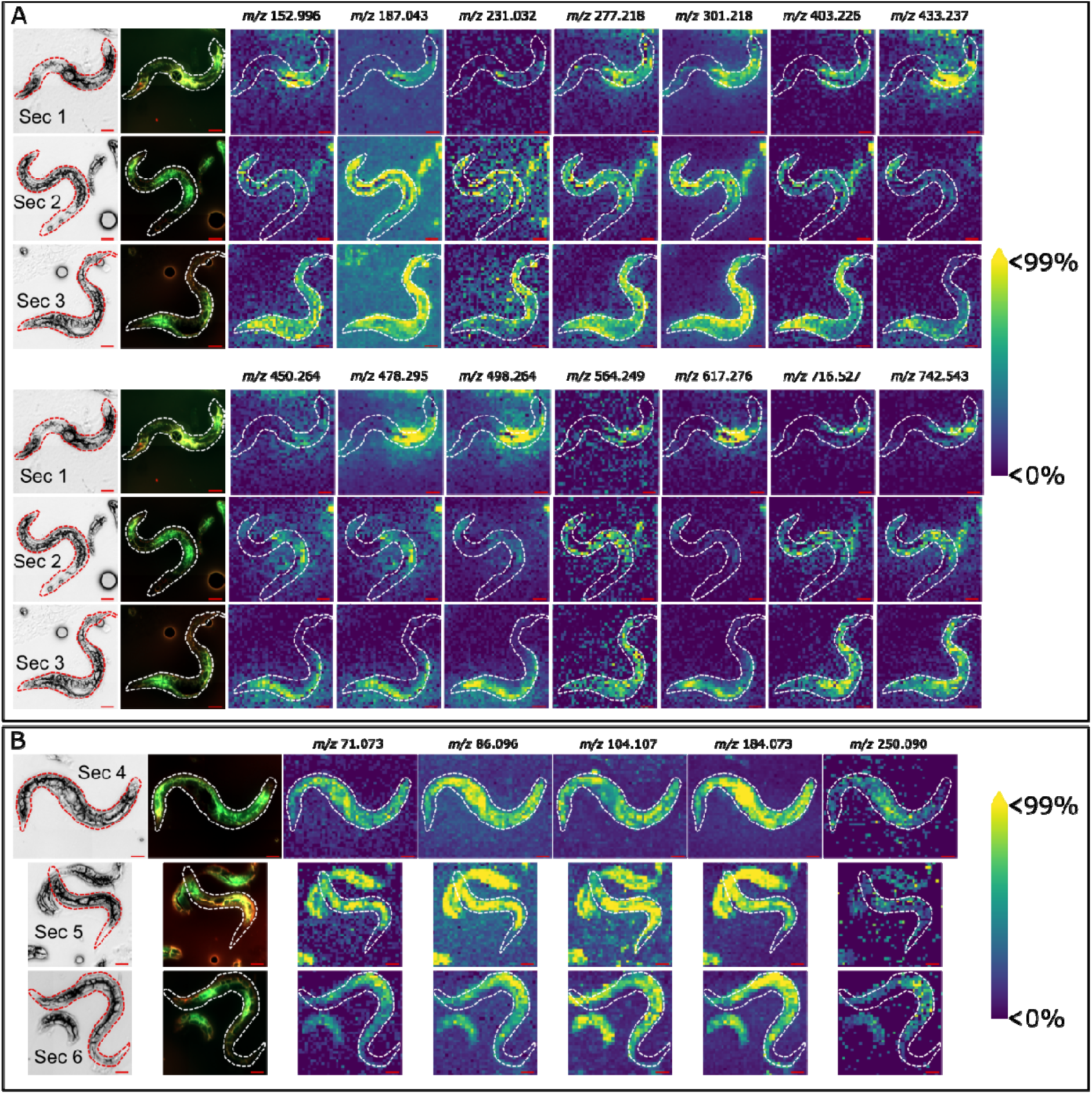
Longitudinal sections of *C. elegans* collected by sandwiching worms in 9:1 FG:PG between blocks of 4:1 PG:FG. Different sections are numbered as Sec 1-6. While a few of these worms (Sec1 and 6) show red fluorescence from clusters of neurons, most sections are longitudinal only through the body of the worm. Both ion images from negative mode MSI (A) and positive mode MSI (B) are presented. For each row, from right to left are brightfield, merged GFP+RFP fluorescence, and ion images of different *m/z*. Scale bar represents 50 μm.

Ion images of three worm sections from negative mode MSI (**Figure 5A**) and positive mode MSI (**Figure 5B**) are shown as examples. Like unoriented sections, we observed that most of the ions are localized within the tissue edges, indicating preservation of ion localizations in worm tissues. Similar co-localization analysis was done to determine ion signal delocalization (**Figure S6**), and results showed that all ions showed high or fair co-localization, indicating that the ions are generally localized within worm tissues (**Table S3**). It should be noted that only the intact worms in the frame were outlined and evaluated, while other fragmented sections in the same frame were not considered for interpretation. Various distribution patterns of ions were observed. Putative annotations based on *m/z* of shown ion images are listed in **Table S2**, while it should be noted that these annotations are only based on MS1 with limited confidence of identification. From negative mode MSI, several ions clearly localized with the GFP signals in the intestinal cells, including *m/z* 433.237, 450.264, 478.295, 498.264, and 617.276 (**Figure 5A**). Interestingly, although all intestine-localized, there were subtle differences in the spatial patterns of ions. For example, only *m/z* of 478.295 was found of relatively high intensity across Section 1 through 3 in **Figure 5A**; however, *m/z* 433.237, 498.264, and 617.276 only showed high intensity in Section 1 and 3, while m/z 450.264 showed highlighted patterns in Section 2 and 3. Since the longitudinal sections can capture anterior and posterior intestine in the same section, such distribution differences can potentially be from the different metabolism in different parts of the intestine. Other distribution patterns include potentially gonad-localized signals (*m/z* 564.249, 716.527, and 742.543) that localize outside intestine and peripheral tissues (*m/z* 187.043, 231.032). Ion signals discovered from positive mode MSI showed less localized distribution compared to what was found in negative mode MSI (**Figure 5B**). PC headgroup precursor (*m/z* 184.073) and fragments (*m/z* 86.096, 71.073) showed co-localization with each other and distribution across worm body. Choline (*m/z* 104.107) showed some patterns along the worm body, however, no clear localization to fluorescence signals were registered.

## Discussion

Our research focused on overcoming challenges in acquiring quality and biological interpretable MSI data for *C. elegans*, a small organism with thick cuticles, via adding another imaging modality of tissue-specific fluorescence reporters expressed in *C. elegans* as well as developing embedding media and oriented sectioning to improve sample preparation for structural integrity. Being able to determine the tissue type and structure is very important for MS imaging data interpretation. Common strategies include pre-MSI autofluorescence imaging and post-MSI staining strategies, however, for *C. elegans*, autofluorescence is limited to intestinal granules and histological stains were rarely reported and unconventional on *C. elegans*.^37,38^ We decided to utilize one of the well-developed toolkits in *C. elegans* research, the genetically encoded fluorescence reporters, to assist the MSI data interpretation. Results showed that the use of fluorescence reporters greatly enhanced our capability to interpret the spatial localization of features mapped to specific worm tissues. We found that the intestinal cytosol GFP was easily identifiable, while the nuclei-localized RFP in neurons suffered from non-specific fluorescence interference but also provided clear, localized signals when detected. In addition, we noticed that the developmental speed was slightly delayed for strains with RFP-tagged neurons for unknown reasons. Future work will explore other multicolored *C. elegans* strains that enable identification of multiple cell types,^39^ reduce non-specific fluorescence by using stronger reporters, and assess the impact of fluorescence tags to worm metabolism.

The optimized gelatin embedding medium composed of both mammalian (porcine) gelatin and fish gelatin leverages the higher gel strength of the porcine gelatin to limit damage to worm bodies while lowering the gelling temperature with the fish gelatin to avoid heat shocking worms. Compared with single-component embedding media, this binary mixture of two different gelatins provided another dimension in tuning the properties of embedding media, making it a recipe of high adaptability when optimizing media for unconventional sample types such as *C. elegans*. The sample preparation process, including embedding and freezing, was simple and fast, minimizing the manipulation of worms which may disturb worm structure and metabolism during sample preparation. Using the optimized embedding media and proper freezing, we were able to generate worm sections with better defined and preserved structures compared to previous reports which is the foundation for ion image interpretation.^17–19^

The sandwiching approach to worm orientation confers the same advantages due to its use of the same embedding medium and furthermore, enables longitudinally oriented sections to be collected consistently with minimal preparation and additional equipment needed. Sandwiching approaches have been a strategy for positioning and sectioning small-sized samples such as *Drosophila* heads and spheroids.^40,41^ In some methods, samples were transferred onto a frozen layer of embedding media, followed by addition of chilled embedding media on the top. However, the temperature differences between frozen layer and newly added media layer could result in freeze-thaw effect for the samples sandwiched in between. In our method, the sandwiching blocks and the in-between embedding media plus worms were assembled at room temperature and then frozen together, avoiding freeze-thaw effect. The addition of dyes to the sandwiching support blocks provided a visual aid to orient the cryostat head for longitudinal sectioning of sandwiched worms. Our approach provides a simple way of oriented sectioning that can be potentially applied to other types of small samples. We noticed that overlapping sections can happen, such as Section 5 (**Figure 5B**), which may confound the spatial interpretation of features. Overlapping sections are difficult to fully get rid of because the worms are randomly distributed in the sandwiching layer and they are still alive and crawling during the sandwiching process. Chances of overlapping worms can be reduced by lowering worm density or manually spreading the worms before putting the top gelatin block to make the sandwiching layer. With sufficient worm populations, we found there are usually sufficient individual sections that can be used for data analysis.

## Conclusion and Limitations

In this work, we focused on overcoming challenges in sample preparation of *C. elegans* for MSI. We have successfully applied tissue-specific fluorescence reporters in *C. elegans* as a new imaging modality on the same tissue to assist spatial assignment of MSI features, optimized binary mixture of fish and porcine gelatin as a new embedding media recipe to achieve preservation of tissue morphology, and developed a liver-worm sandwiching strategy for embedding to consistently split the *C. elegans* in the longitudinal sections. By achieving structural preservation and enabling fluorescence reporters as an additional imaging modality for data interpretation, this workflow overcomes key limitations in MSI of *C. elegans* and provides a practical framework for MSI analysis of microscale biological samples.

Meanwhile, we identify limitations for future improvement. First, while we consistently obtain longitudinal sections from the sandwiching method, it was still very challenging to obtain a perfect nose-to-tail longitudinal section. This resulted from multiple factors. For example, the softness of gelatin sandwiching blocks as well as the presence of meniscus at their edge made it possible to have an uneven surface for the worms embedded in between. Additionally, even with the guidance of block colors, the angle of cryostat blade can still be slightly off. Future improvements can focus on creating flat space in between sandwiching blocks, finding ways to calculate and fine-tune the blade angle and image serial sections to obtain different layers of the longitudinal section (**Figure S8**). The second limitation is related to the size of *C. elegans*. With a maximal width below 100 μm, it is challenging to resolve single cells in their body with our current pixel resolution at 10-μm. Expansion MSI has recently been developed to show significantly enhanced spatial resolution on biological samples.^42–44^ While the thick cuticle in *C. elegans* represents a challenge to achieve tissue expansion, it still presents an opportunity for future high-resolution MSI in *C. elegans*. Thirdly, analyte delocalization could result from sample preparation and/or small molecule being secreted or diffused into the embedding media, particularly as we embedded the *C. elegans* alive to avoid freeze-thaw cycles. With the observation of some delocalization patterns, drier matrix application techniques such as sublimation should be explored to reduce technical-sourced delocalization. Fourth, current spatial assignment of features relied on image overlapping and visual inspection. Future work should explore image co-registration strategies between MSI and fluorescence microscopy for more robust spatial assignment of tissues for biological conclusions. Lastly, the metabolite annotation presented a challenge to us during this work. We searched for the precursor *m/z* against Metabolomics Workbench, Human Metabolome Database, and LIPID MAPS, as well as a manually curated, *C. elegans*-specific metabolome database.^23^ However, putative annotations solely based on MS1 have low confidence of identification. Future work will include performing on-tissue MS/MS with the iprm-PASEF mode available in timsTOF fleX, incorporating trapped ion mobility spectrometry (TIMS) for collision cross section (CCS) matching, and adding bulk LC-MS metabolomics and lipidomics to build library and assist metabolite annotation.

## Supporting information

WormSandwiching movie

## Associated Content

### Supporting Information

Average mass spectra of *C. elegans* sections, mass resolution, brightfield and fluorescence images of sectioned and intact worms, *m/z* putative annotations, and co-localization analysis.

### Data Availability Statement

All raw MSI data and method files are uploaded to Mass Spectrometry Interactive Virtual Environment (MassIVE; massive.ucsd.edu) under accession ID MSV000102830.

### Author Contributions

R.E.J.: conceptualization, methodology, validation, formal analysis, investigation, data curation, visualization, writing – original draft; E.W.S.: methodology, investigation, writing – review & editing; Y.S.: methodology, resources, writing – review & editing; T.A.Q.: conceptualization, methodology, resources, data curation, writing – review & editing, supervision, project administration, funding acquisition.

## Acknowledgements

Some *C. elegans* and *E. coli* strains were purchased from the Caenorhabditis Genetics Center (University of Minnesota, Minneapolis, USA), which is funded by the National Institutes of Health, National Center for Research Resources. This work was supported by startup funding from Michigan State University to T.A.Q. This study was conducted during Y.S.’s study abroad as a research fellow, supported by his employment at Kitasato University, which provided continuous salary support. E.W.S. was supported by an REU program awarded to the Department of Chemistry at Michigan State University under NSF CHE-2150173.

## For Table Of Contents Use Only

The figure depicts an abbreviated version of the workflow used to orient, section, and image *C. elegans* nematodes. It shows the process of rinsing gravid adults off a culture plate followed by addition with embedding medium to a solid block of gelatin prior to sandwiching. This is followed by cryosectioning and imaging, with both microscope and MS imaging modalities represented by a drawing of the Bruker timsTOF fleX. The final image depicts brightfield, fluorescence, and MS images of an example nematode.

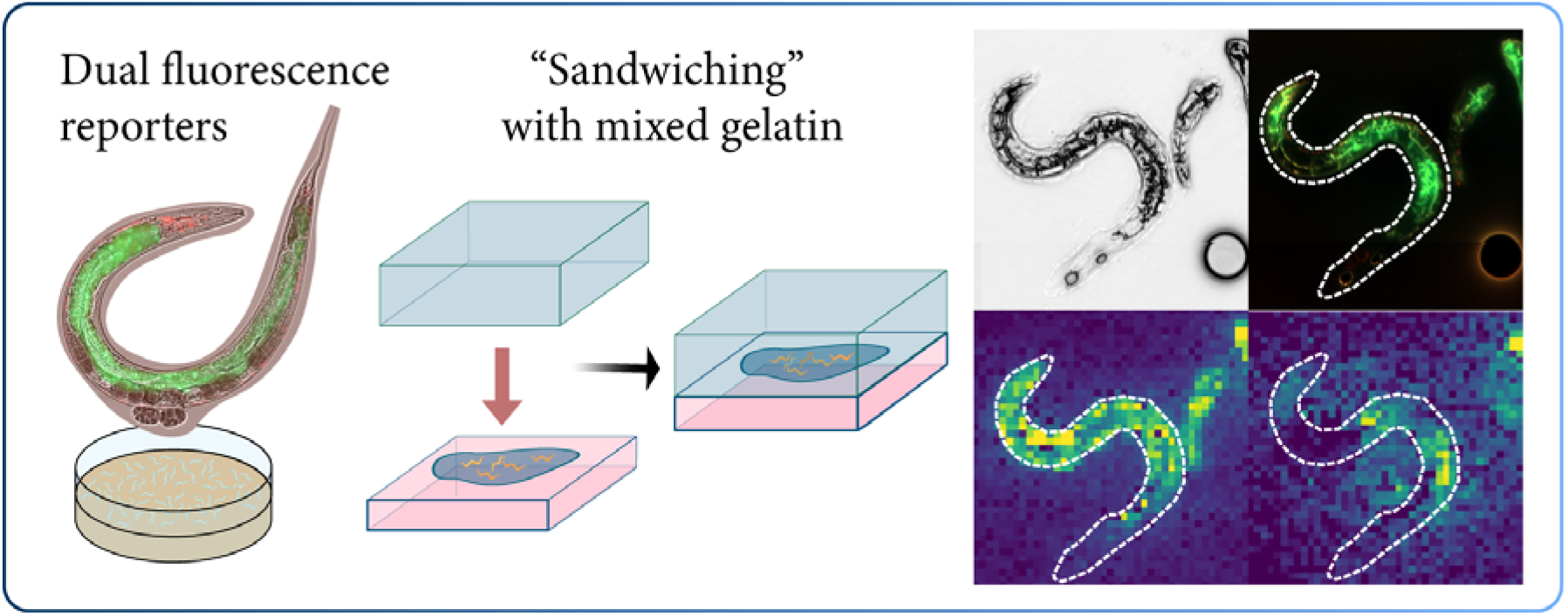

