## Supplemental Information for "Mass Spectrometry Imaging of *Caenorhabditis elegans* with Fluorescence Microscopy of Tissue-Specific Reporters and Oriented Cryo-Sectioning"

**Table of Contents**

**Table S1**. Mass resolution for select positive (CHCA) and negative (DAN) mode chemical matrix ions.

**Figure S1**. Microscopic images of the cross sections from unoriented embedding with different combinations of embedding media and freezing methods.

**Figure S2**. Microscopic images of the longitudinal sections from unoriented embedding with different combinations of embedding media and freezing methods.

**Figure S3**. Whole-worm image of live dual-reporter worm strain.

**Figure S4**. Average spectra of *C. elegans* and embedding media for positive mode with CHCA.

**Figure S5**. Average spectra of *C. elegans* and embedding media for negative mode with DAN.

**Table S2**. Putative annotations and mass accuracy for select metabolite and matrix ion signals.

**Figure S6**. Example overview of longitudinal sections from sandwiching.

**Table S3.** Correlation values for features shown in **Figure 3** and **Figure** **5**.

**Figure S7**. An example of zoomed-out overview of longitudinal sections from sandwiching.

**Figure S8**. Zoomed in images of neurons in head ganglia and VNC.

**Figure S9**. Brightfield and fluorescence images of adjacent sections of oriented worms.

**Table S1**. Mass resolution for selected positive (CHCA) and negative (DAN) mode chemical matrix ions. Twenty spectra for each matrix were randomly sampled and the resolutions for each *m/z* averaged across them. The average resolution shown is that of all *m/z* for each chemical matrix.

| **DAN ions** | **156.0692** | **157.0775** | **313.1455** | **471.1455** | **Average** |
| --- | --- | --- | --- | --- | --- |
| **Resolution** | 24058 | 21251 | 26642 | 36663 | 27153 |
| **CHCA ions** | **190.0499** | **212.0324** | **221.9718** |  | **Average** |
| **Resolution** | 28414 | 26513 | 25405 |  | 26773 |

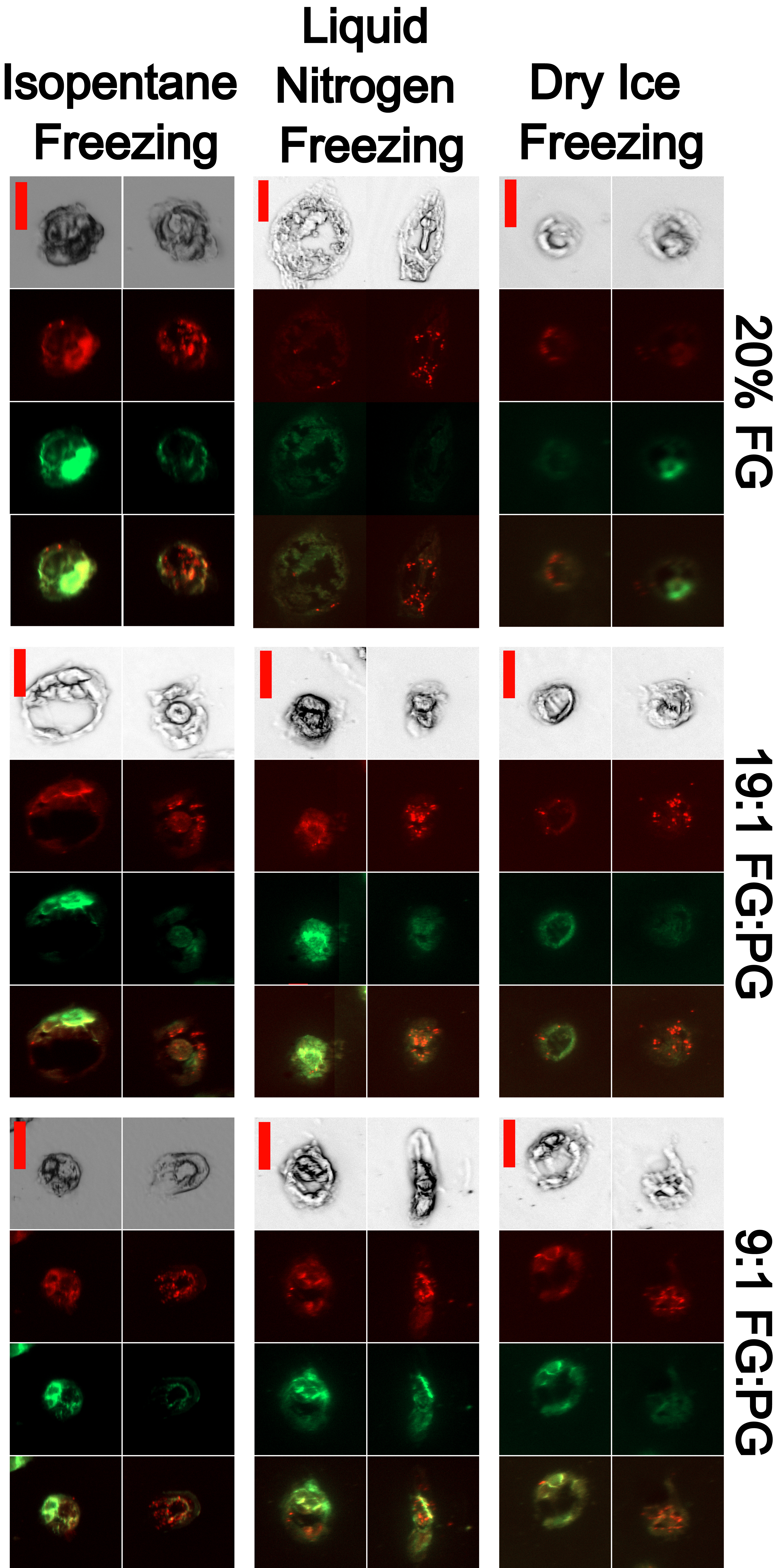

**Figure S1**. Cross sections from unoriented embedding with different combinations of embedding media and freezing methods. In each sub-panel, the columns from left to right represent: brightfield, RFP, GFP and merged fluorescence image. Scale bar represents 50 μm.

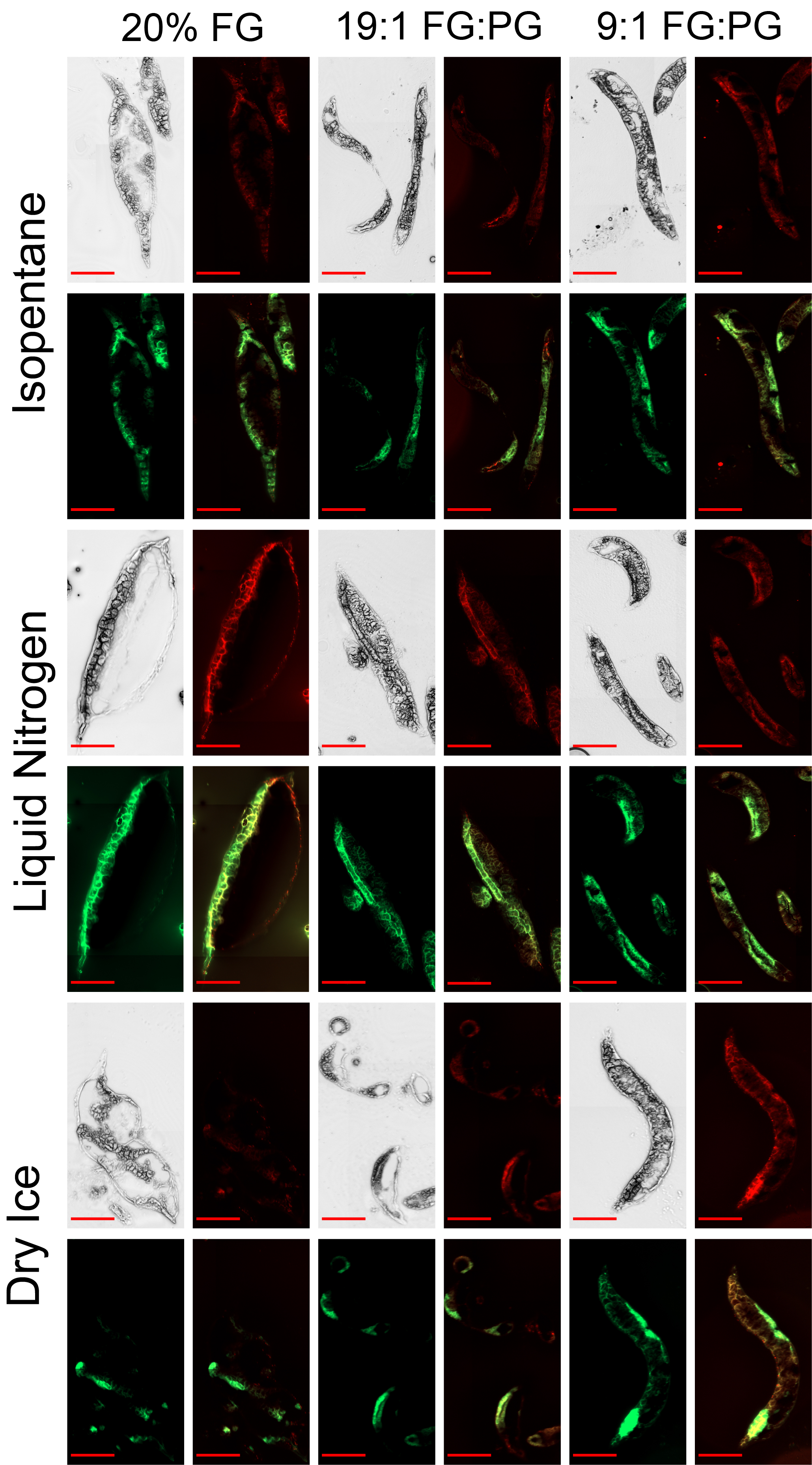
 **Figure S2**. Longitudinal sections from unoriented embedding with different combinations of embedding media and freezing methods. In each sub-panel, the columns from left to right represent: brightfield, RFP, GFP and merged fluorescence image. Scale bar represents 50 μm.

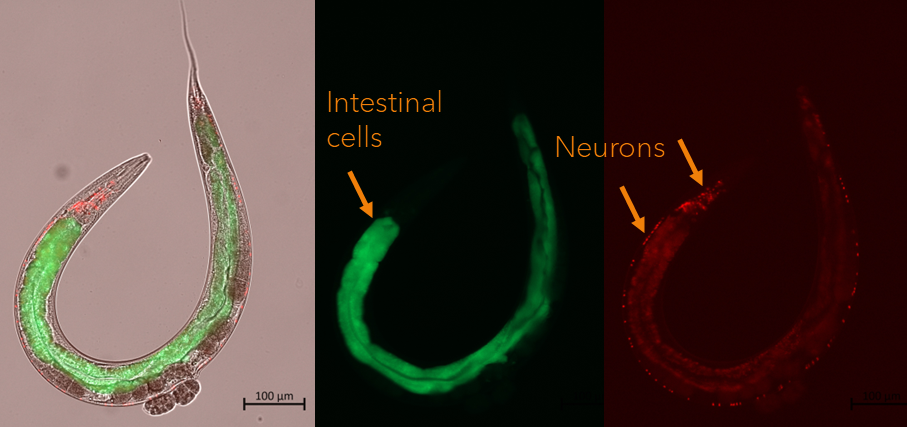

**Figure S3**. Whole-worm image of live dual-reporter strain. Left: merged images of brightfield, GFP, and RFP. Middle: GFP. Right: RFP.

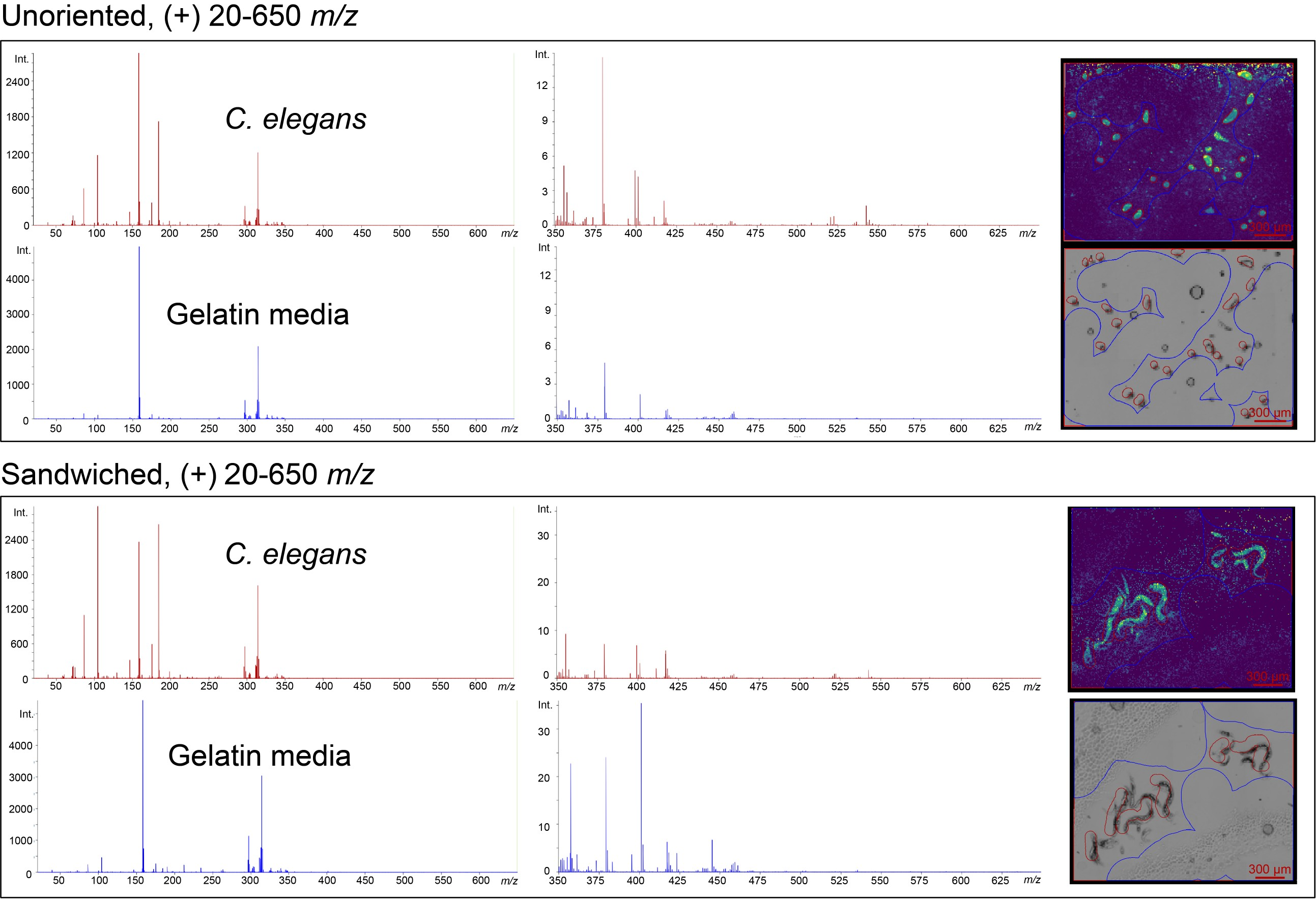

**Figure S4**. Average spectra for different regions of *C. elegans* sections using positive mode analysis with CHCA as the MALDI matrix. Left column: average spectra of full detection *m/z* range; middle column: a zoomed-in spectra of higher m/z; right column: illustration of drawn regions to acquire the average spectra (ion image on top and optical image at bottom). Regions were hand-drawn based on ion images to cover *C. elegans* sections. Worm regions were drawn slightly larger than ion images to ensure that any worm signals associated with the cuticle were included in the average spectra. Red regions represent worms, with blue regions drawn over areas of 9:1 FG:PG which contain no worms. Note that the regions drawn based on ion images did not fully align with optical images and show offsets of ~50 μm, which is likely because of the minor errors in setting up teaching points.

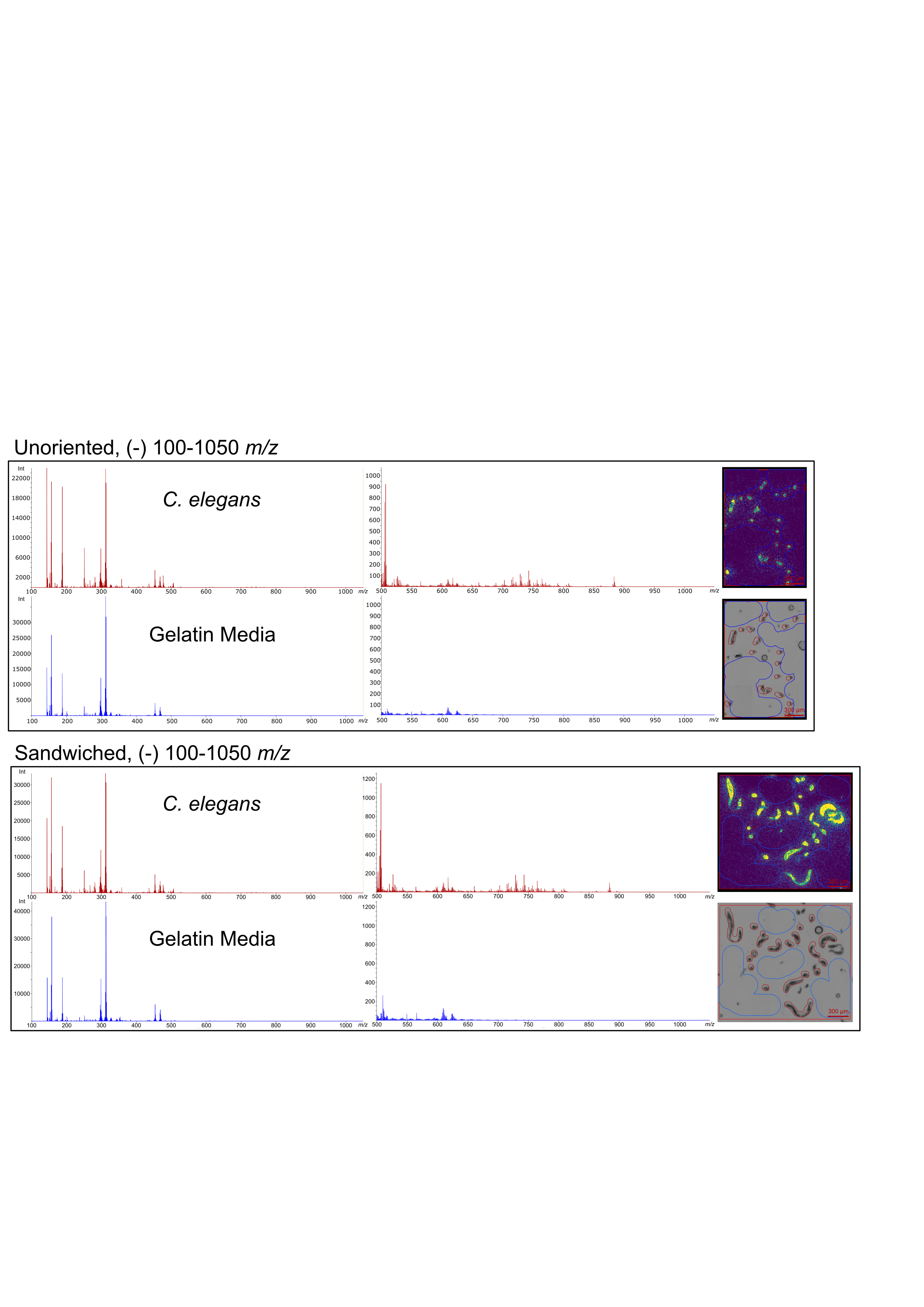
**Figure S5**. Average spectra for different regions of *C. elegans* sections using negative mode analysis with DAN as the MALDI matrix. Regions were hand-drawn to be representative rather than covering every pixel of the section as this is sufficient for generating average spectra. Worm regions were drawn slightly larger than ion images to ensure that any worm signals associated with the cuticle were included in the average spectra. Red regions represent worms, with blue regions drawn over areas of 9:1 FG:PG which contain no worms. The second column of average spectrum focuses on the high *m/z* range of each spectrum for better visibility.

**Table S2**. Putative annotations based on MS1 of features identified from MSI runs.

| Features from unoriented and sandwiched imaging run, CHCA, (+) 20-650 *m/z* | | | | | |
| --- | --- | --- | --- | --- | --- |
| **Feature *m/z*** | **Annotation based on MS1 accurate mass** | **Molecular Formula** | **Adduct** | **Adduct Mass** | **Delta (ppm)** |
| 71.073 | PC fragment | C4H9N | [M]^+^ | 71.073 | 6 |
| 86.096 | PC fragment | C5H11N | [M]^+^ | 86.097 | 12 |
| 104.107 | choline | C5H14NO | [M]^+^ | 104.108 | 5 |
| 146.117 | acetylcholine | C7H16NO2 | [M]^+^ | 146.118 | 8 |
| 184.073 | phosphorylcholine | C5H15NO4P | [M+H]^+^ | 184.074 | 5 |
| 250.090 | S-acetyldihydrolipoamide | C10H19NO2S2 | [M+H]^+^ | 250.090 | 0 |
| Features from unoriented and sandwiched imaging run, DAN, (-) 100-1050 *m/z* | | | | | |
| **Feature *m/z*** | **Annotation based on MS1 accurate mass** | **Molecular Formula** | **Adduct** | **Adduct Mass** | **Delta (ppm)** |
| 152.996 | 2-Phosphonopropionic acid | C3H7O5P | [M-H]^-^ | 152.996 | 2 |
| *187.043* | *[No match]* |  |  |  |  |
| 231.032 | *[No match]* |  |  |  |  |
| 257.228 | *[No match]* |  |  |  |  |
| 267.234 | FA 17:1 | C17H32O2 | [M-H]^-^ | 267.233 | 2 |
| 277.218 | FA 18:3 | C18H30O2 | [M-H]^-^ | 277.217 | 3 |
| *299.054* | *[No match]* |  |  |  |  |
| 301.218 | FA 20:5 | C20H30O2 | [M-H]^-^ | 301.217 | 2 |
| 304.237 | *[No match]* |  |  |  |  |
| *315.033* | *[No match]* |  |  |  |  |
| 403.226 | CPA 17:1 | C20H37O6P | [M-H]^-^ | 403.226 | 2 |
| 417.242 | LPA O-18:3 | C21H39O6P | [M-H]^-^ | 417.241 | 2 |
| 431.258 | LPA O-19:3 | C22H41O6P | [M-H]^-^ | 431.257 | 2 |
| 433.237 | LPA 18:2 | C21H39O7P | [M-H]^-^ | 433.236 | 2 |
| 450.264 | LPE 16:1 | C21H42NO7P | [M-H]^-^ | 450.263 | 3 |
| *477.108* | *[No match]* |  |  |  |  |
| 478.295 | LPE 18:1 | C23H46NO7P | [M-H]^-^ | 478.294 | 3 |
| 498.264 | LPE 20:5 | C25H42NO7P | [M-H]^-^ | 498.263 | 3 |
| 503.284 | *[No match]* | C26H40N4O6 | [M-H]^-^ | 503.288 | 7 |
| 520.306 | PE 20:1 | C25H48NO8P | [M-H]^-^ | 520.305 | 3 |
| *564.249* | *[No match]* |  |  |  |  |
| 617.276 | LPI 20:5 | C29H47O12P | [M-H]^-^ | 617.273 | 4 |
| 702.512 | PE 33:1 | C38H74NO8P | [M-H]^-^ | 702.508 | 5 |
| 714.512 | PE 34:2 | C39H74NO8P | [M-H]^-^ | 714.508 | 5 |
| 716.527 | PE 34:1 | C39H76NO8P | [M-H]^-^ | 716.524 | 4 |
| 721.483 | PA 38:5 | C41H71O8P | [M-H]^-^ | 721.481 | 3 |
| 728.563 | PE O-36:2 | C41H80NO7P | [M-H]^-^ | 728.560 | 4 |
| 730.577 | PE(P-36:0) | C41H82NO7P | [M-H]^-^ | 730.576 | 2 |
| 742.543 | PE 36:2 | C41H78NO8P | [M-H]^-^ | 742.539 | 4 |
| 758.575 | PE 37:1 | C42H82NO8P | [M-H]^-^ | 758.571 | 6 |
| 776.527 | PE 39:6 | C44H76NO8P | [M-H]^-^ | 776.524 | 4 |
| 883.537 | PI 38:5 | C47H81NO13P | [M-H]^-^ | 883.534 | 3 |
| 884.543 | PS 44:9 | C50H80NO10P | [M-H]^-^ | 884.545 | 2 |

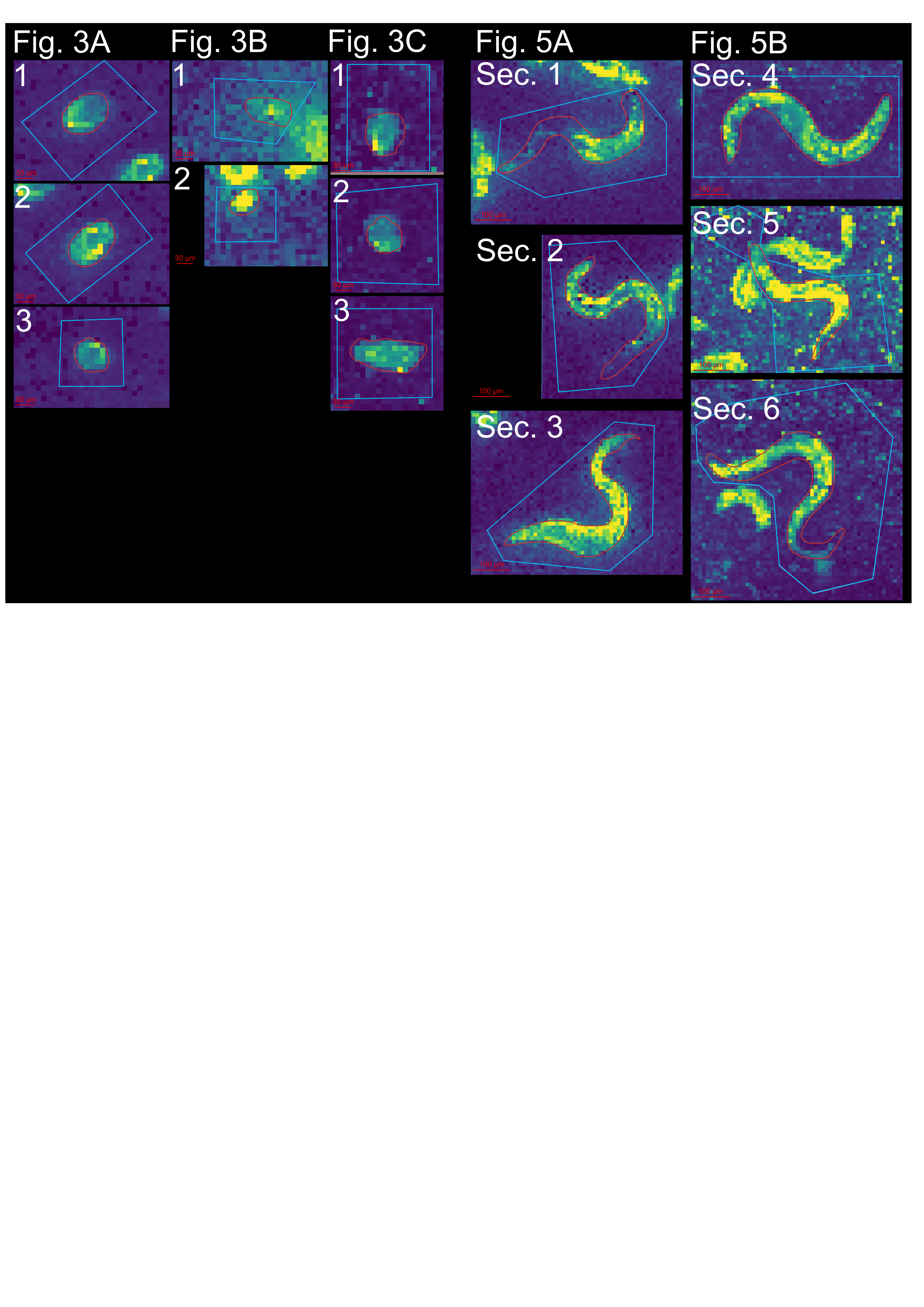

**Figure S6.** Mass spectrometry images showing the regions used to calculate correlation values in SCiLS to determine the degree of analyte delocalization. One ROI was drawn around the worm tissue edge, and another was drawn around the worm section while avoiding other nearby worm tissues. Images correspond to the figures and sections as labeled, with images for **Figure 3** numbered with 1 as the top section in that figure. Scale bars are 30 μm for Figure 3 images and 100 μm for Figure 5. Ion images used to show ROIs are the same for all sections in a given figure panel and are as follows: *m/z* 187.043 for Fig. 3A, 267.234 for Fig. 3B, 104.107 for Fig. 3C, 301.218 for Fig. 5A, and 104.107 for Fig 5B.

**Table S3.** Correlation values for features shown in **Figure 3** and **Figure** **5**. Regions were chosen as shown in **Figure S9**. Values below 0.2 (yellow) are considered to have poor co-localization, while those above 0.5 (blue) show high co-localization with the worm tissue region. All other values indicate fair co-localization and are highlighted in green.

| **FIGURE 3A** | | | | **FIGURE 5A** | | | |
| --- | --- | --- | --- | --- | --- | --- | --- |
| *m/z* | SECTION 1 | SECTION 2 | SECTION 3 | *m/z* | SECTION 1 | SECTION 2 | SECTION 3 |
| 187.043 | 0.854 | 0.882 | 0.840 | 152.996 | 0.422 | 0.519 | 0.756 |
| 257.228 | 0.847 | 0.742 | 0.728 | 187.043 | 0.443 | 0.582 | 0.777 |
| 477.108 | 0.582 | 0.568 | 0.645 | 231.032 | 0.268 | 0.338 | 0.484 |
| 520.306 | 0.617 | 0.624 | 0.631 | 277.218 | 0.505 | 0.645 | 0.784 |
| 702.512 | 0.791 | 0.735 | 0.401 | 301.218 | 0.631 | 0.728 | 0.847 |
| 714.512 | 0.819 | 0.728 | 0.638 | 403.226 | 0.491 | 0.596 | 0.763 |
| 758.575 | 0.742 | 0.672 | 0.645 | 433.267 | 0.359 | 0.394 | 0.617 |
| 776.527 | 0.659 | 0.638 | 0.652 | 450.264 | 0.213 | 0.345 | 0.429 |
| 884.543 | 0.700 | 0.645 | 0.666 | 478.295 | 0.373 | 0.561 | 0.638 |
| **FIGURE 3B** | | | | 498.264 | 0.359 | 0.394 | 0.547 |
| *m/z* | SECTION 1 | SECTION 2 |  | 564.249 | 0.289 | 0.429 | 0.575 |
| 257.228 | 0.512 | 0.653 |  | 617.276 | 0.394 | 0.310 | 0.603 |
| 267.234 | 0.638 | 0.728 |  | 716.527 | 0.352 | 0.463 | 0.686 |
| 299.055 | 0.463 | 0.629 |  | 742.543 | 0.394 | 0.526 | 0.756 |
| 304.237 | 0.409 | 0.772 |  | **Figure 5B** | | | |
| 315.033 | 0.654 | 0.386 |  | *m/z* | SECTION 4 | SECTION 5 | SECTION 6 |
| 417.242 | 0.580 | 0.680 |  | 71.073 | 0.798 | 0.714 | 0.693 |
| 431.258 | 0.556 | 0.675 |  | 86.096 | 0.840 | 0.721 | 0.700 |
| 503.284 | 0.639 | 0.153 |  | 104.107 | 0.819 | 0.700 | 0.672 |
| 728.564 | 0.549 | 0.599 |  | 184.073 | 0.833 | 0.798 | 0.742 |
| **FIGURE 3C** | | | | 250.090 | 0.603 | 0.436 | 0.519 |
| *m/z* | SECTION 1 | SECTION 2 | SECTION 3 |  |  |  |  |
| 71.073 | 0.777 | 0.861 | 0.829 |  |  |  |  |
| 86.096 | 0.777 | 0.875 | 0.839 |  |  |  |  |
| 104.107 | 0.735 | 0.854 | 0.859 |  |  |  |  |
| 146.117 | 0.350 | 0.429 | 0.739 |  |  |  |  |
| 184.073 | 0.833 | 0.930 | 0.879 |  |  |  |  |

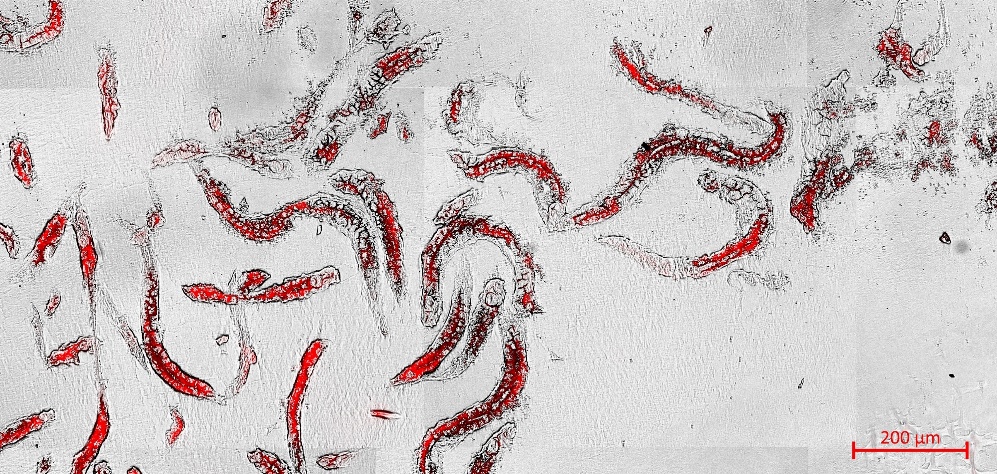

**Figure S7**. An example of zoomed-out overview of longitudinal sections from sandwiching.

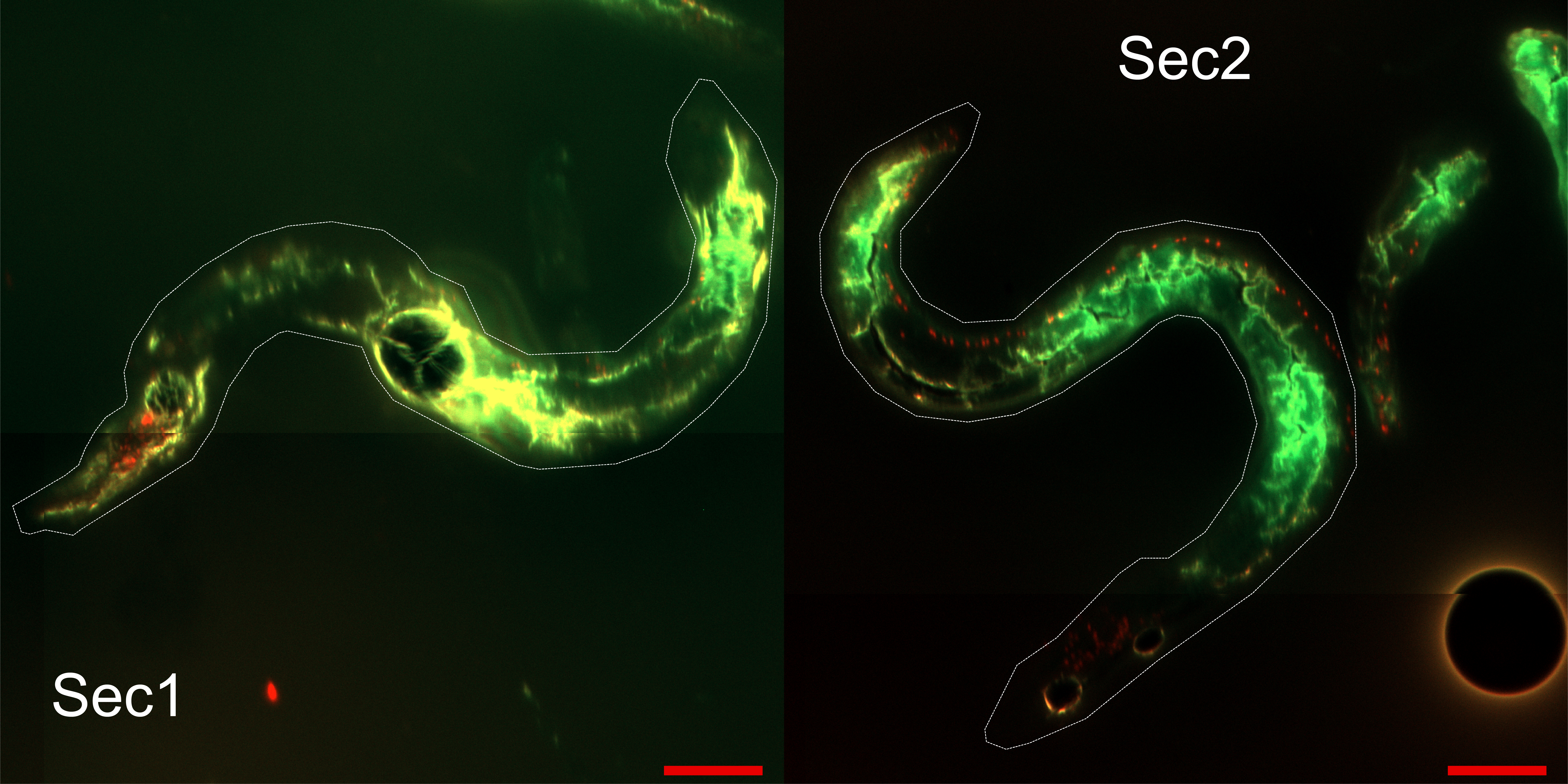

**Figure S8**. Zoomed in images of neurons in Section 1 and Section 2. In Section 1 (left), RFP from head ganglia can be clearly observed. In Section 2 (right), RFP from VNC neurons along one side of the worm body can be observed.

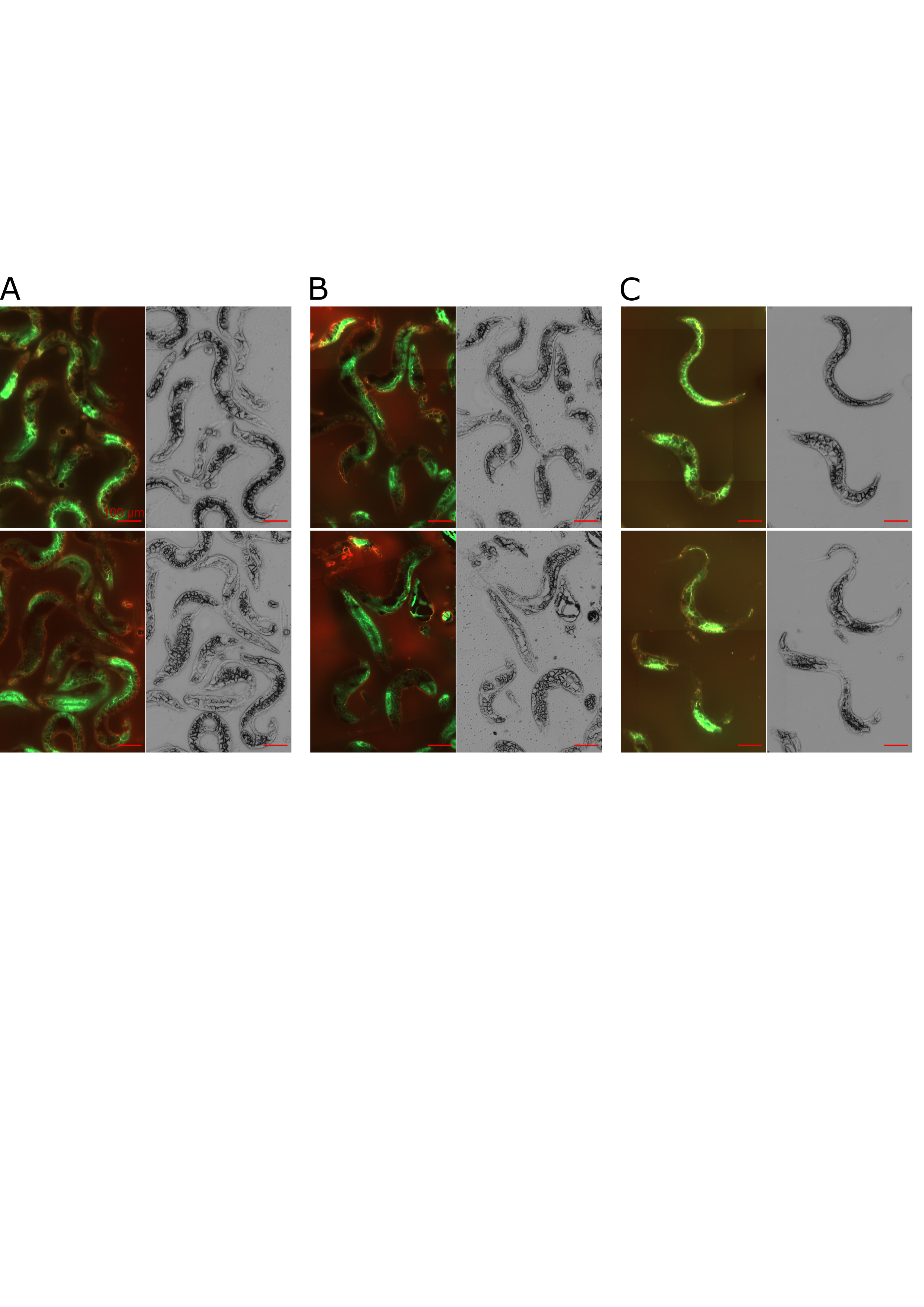

**Figure S9**. Brightfield and fluorescence images for three pairs of adjacent sections from sandwiched sample preparation. Panels A, B, and C each contain the brightfield and fluorescence images for one section in the top row and those for its adjacent section in the bottom row. Scale bars represent 100 μm.
